# Loss of *UBP1* drives oxaliplatin resistance through a targetable dependency on translation initiation

**DOI:** 10.64898/2026.05.18.725866

**Authors:** Moustafa Abohawya, Tim Schmäche, Julia Dietzel, Ina Hollerer, Li Ding, Maciej Paszkowski-Rogacz, Rico Barsacchi, Therese Seidlitz, Sebastián García Tobar, Frank Buchholz, Daniel Stange, Jovan Mirčetić

## Abstract

Oxaliplatin is a common component of various chemotherapeutic regimens for the treatment of gastrointestinal cancers. However, the majority of patients exhibit resistance to oxaliplatin-based therapy. Here, we integrated knockout and transcription-activation CRISPR screens in patient-derived gastric cancer organoids (GC PDOs) to comprehensively profile major genetic and transcriptomic changes observed over the course of resistance acquisition. Our screens identified UBP1, a transcription factor frequently lost in GC, as a critical determinant of oxaliplatin resistance development. Leveraging a large GC organoid biobank from a co-clinical trial and primary tumor omics data, we reveal that downregulation of specifically MYC-driven ribosome biogenesis drives oxaliplatin resistance, highlighting the drug’s role as a ribosome biogenesis stressor. Mechanistically, *UBP1* loss reduced expression of its direct target, MAX, a MYC cofactor, leading to downregulation of ribosome biogenesis and protection against nucleolar stress. Crucially, we discover that such downregulation is inevitably followed by a compensatory reliance on translation initiation, making it a therapeutic vulnerability in oxaliplatin-resistant tumors. Consequently, the resistance could be overcome by a synergistic action of the translation initiation repressor 4EGI, and the effect was also maintained in PDO that acquired resistance *in vivo* under clinically relevant conditions. Our data uncover a common marker of oxaliplatin resistance and identify a novel therapeutic strategy to reverse resistance to one of the most frequently used anticancer drugs.

## Introduction

Gastric cancer (GC) is the fifth most common malignancy worldwide^1^. A recent breakthrough in its treatment emerged with the introduction of the FLOT regimen (5-fluorouracil, oxaliplatin, docetaxel, and leucovorin), which showed improved overall survival compared with the previously established ECF regimen^2^. Two changes distinguish FLOT from ECF: the replacement of cisplatin with oxaliplatin and epirubicin with docetaxel. FLOT has, hence, become the standard of care in gastric cancer.

Oxaliplatin, a next-generation platinum compound, was initially approved for colorectal cancer, demonstrating substantial therapeutic efficacy^3^. Now, oxaliplatin has become a cornerstone of treatment for a range of gastrointestinal malignancies, including colon^4^, gastric^5^, and esophageal cancers^6^. Notably, even in the context of FLOT efficacy in GC, response rates remain suboptimal: approximately 37% of patients achieve a major response, while the majority exhibit a partial or no response^2,7^. Therefore, a significant proportion of patients show resistance to FLOT, underscoring the need to define the determinants of its therapeutic action, including those of oxaliplatin.

The mechanistic explications of oxaliplatin’s action remain largely controversial. New evidence points to its ability to induce stress in the biogenesis and/or function of the translation machinery, differentiating it from some of the other platinum compounds that induce apoptosis via the DNA damage response^8–10^. These findings provided a potential explanation for the clinical utility of oxaliplatin in colorectal cancer, which is characterized by high rates of protein translation^9^. However, the clinical relevance of such observations, as well as the perspective they offer on the problem of therapeutic resistance, remains unclear.

In this study, we set out to identify the genetic determinants of oxaliplatin resistance in gastric cancer. Through a pair of comprehensive loss- and gain-of-function CRISPR screens in patient-derived organoids (PDOs), we uncover that a frequent loss of transcription factor UBP1 causes oxaliplatin resistance in GC by downregulating MYC-driven ribosome biogenesis. Crucially, this downregulation is compensated for by an increase in translation initiation rates, rendering the resistant cells vulnerable to translation blockers. Our work, therefore, identifies a common genetic marker of oxaliplatin resistance and suggests an alternative treatment approach potentially applicable to one-third of all GC patients.

## Results

### Forward-genetics pipeline in GC PDOs reveals resistance drivers

To delineate oxaliplatin resistance drivers, we first aimed to model the evolution of gastric cancer resistance acquisition. We used a patient-derived organoid line (hereafter, DD109 parental) that was sensitive to oxaliplatin. DD109 was established from a chromosomally unstable (CIN) gastric cancer^11^, the most common GC subtype. Successive rounds of treatment (10 cycles) led to the development of an oxaliplatin-selected PDO line (hereafter, oxa-selected) that was 16 times more resistant than its parental counterpart (Figure 1A).

**Figure 1.**
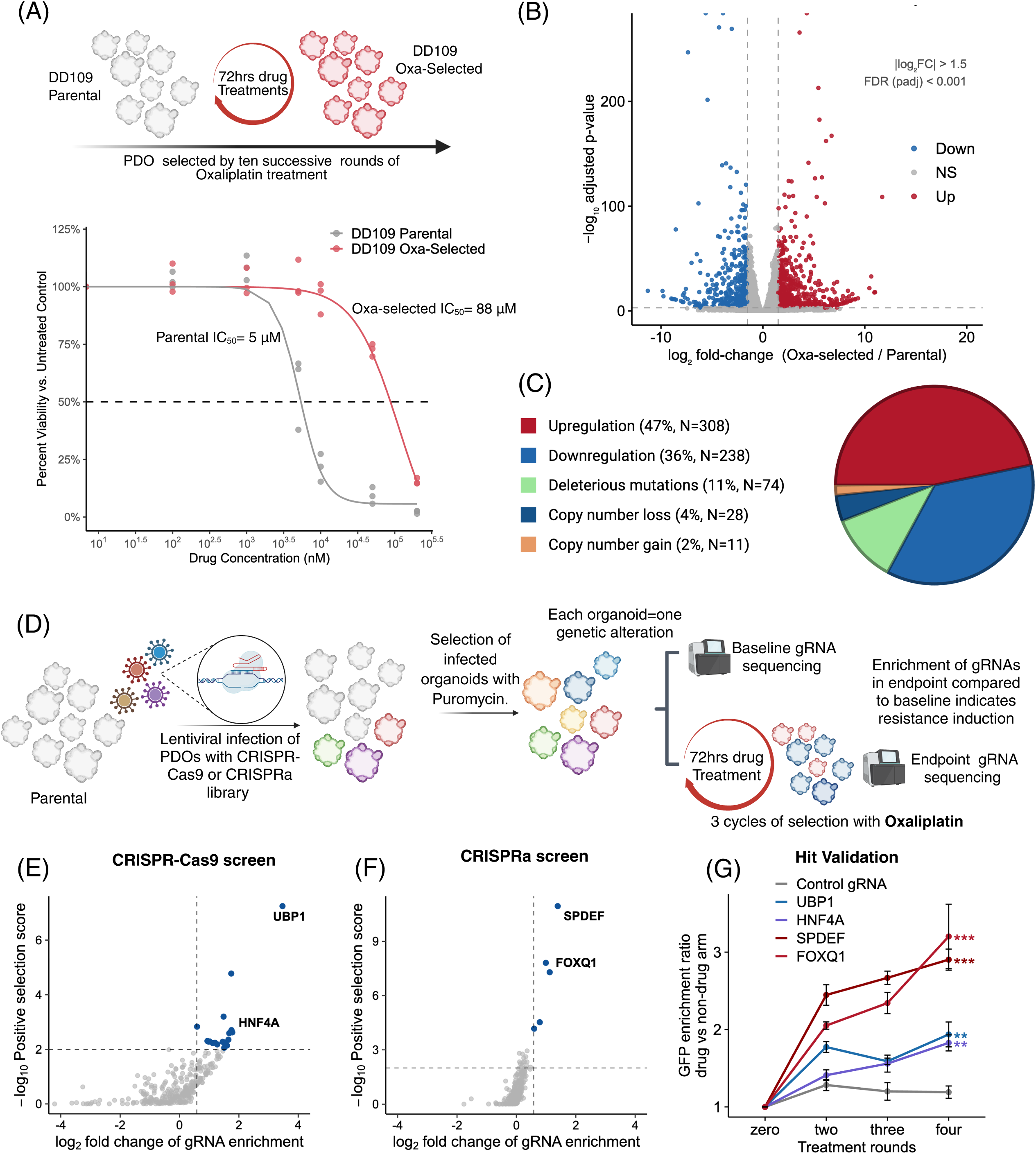
Forward□genetics pipeline using patient□derived gastric cancer organoids uncovers drivers of oxaliplatin resistance. **(A)** Generation of oxaliplatin□resistant DD109 PDOs by subjecting parental organoids to successive 72 h cycles of 5-10 μM oxaliplatin (top schematic). Dose-response viability curves for parental (n=3) versus oxaliplatin-selected DD109 PDOs (n=3; bottom); percent viability is normalized to untreated control. (B) Volcano plot of differential expression in oxaliplatin-selected versus parental DD109 PDOs. Red and blue points denote genes meeting |log□□FC| > 1.5 and FDR < 0.001 (dashed lines). (C) Distribution of “functionally altered” genes between resistant and parental PDOs. (D) Schematic of CRISPR□Cas9 knockout and CRISPR activation (CRISPRa) screens in DD109 PDOs. (E, F) Volcano plots from the CRISPR-Cas9 knockout (E) and CRISPRa (F) screens showing log□ fold-change (FC) of sgRNA enrichment versus –log□□ positive□selection score. Dashed lines denote hit thresholds: positive selection score < 0.01 and FC > 1.5. (G) Hit validation by GFP-competition assays. The percentage of GFP+ organoids in the drug arm (n=3) normalized to the non-drug arm was compared between each hit and a targeting control via linear mixed effect model (***p < 0.001, **p < 0.01).

Next, we aimed to identify which genes were functionally altered between the two isogenic organoid lines. RNA-seq analysis revealed hundreds of genes that were differentially expressed (|log_2_FC⎪ > 1.5 and p_adj_ < 0.001, Figure 1B, Supplementary Table S1), indicating a substantial selection pressure exerted on the oxa-selected line. Additionally, we used exome sequencing to identify genes that had acquired functionally deleterious mutations and copy number changes (Supplementary Table S2-3). In total, a list of differentially altered genes between the oxa-selected and the parental DD109 was compiled, comprising 308 upregulated genes, 238 downregulated genes, 74 genes with deleterious mutations, 28 genes with copy number loss, and 11 genes with copy number gain (Figure 1C).

To functionally profile and pinpoint direct drivers of resistance, we turned to a CRISPR-based screening approach we have recently optimized for organoid biology^12^. For genes that had lost their function due to downregulation, copy number loss, or mutation, we utilized CRISPR-Cas9 (Supplementary Table S4). Additionally, we deployed a transcription activation (CRISPRa^13^) modality for genes that were either upregulated or gained a copy number (Supplementary Table S5). In both modalities, a lentiviral library of guide RNAs (gRNAs) targeting the genes of interest was used to infect parental DD109. After infection, a heterogeneous organoid population was obtained, encompassing all significant genetic/transcriptomic changes observed during the acquisition of resistance. Organoids were then subjected to three cycles of oxaliplatin selection, and the guides were sequenced pre- and post-selection to identify those with a resistance-driving capacity (Figure 1D).

The CRISPR-Cas9 screen identified ten potential hits (Figure 1E, Supplementary Table S6), while the CRISPRa screen added five additional potential hits (Figure 1F, Supplementary Table S7, a threshold of positive selection score p_pos.sel_ < 0.01 and FC > 1.5). To validate the hits, we used an organoid-competition assay (Figure S1). In this pipeline, organoids infected with gRNA-GFP for each hit compete with untagged wild-type counterparts and undergo four successive rounds of oxaliplatin treatment. Four genes showed a consistent, progressive GFP enrichment over time, validating them as functionally relevant oxaliplatin-resistance drivers: *UBP1* and *HNF4A* knockouts (KO), and *SPDEF* and *FOXQ1* upregulations (Figure 1E). Of note, all gRNAs were highly efficient in inducing either KO (*UBP1* and *HNF4A*, Figure S2A-B) or gene upregulations (*SPDEF* and *FOXQ1*, Figure S2C-D). Taken together, our comprehensive screens profiled major genetic/transcriptomic changes occurring during resistance acquisition and identified four transcription factors as resistance drivers: loss of *UBP1* and *HNF4A*, and upregulation of *SPDEF* and *FOXQ1*.

### The screen hits are transcriptionally interconnected and drive downregulation of ribosome biogenesis

Since all identified drivers of oxaliplatin resistance were transcription factors, we investigated the transcriptional consequences of their respective knockout or upregulation. Four PDO lines were developed for this purpose – knockout of *UBP1* and *HNF4A* and upregulation of *SPDEF* and *FOXQ1* – using the validated gRNAs from organoid competition assays (Figure 2A). We first investigated whether the factors formed transcriptional networks and mutually regulated each other. Notably, both UBP1 KO and HNF4A KO resulted in upregulation of the other two factors (Figure 2A). The opposite was not true; the upregulation of *SPDEF* or *FOXQ1* did not result in uniform downregulation of *UBP1* and *HNF4A* (Figure S3A-B). Therefore, loss of either *UBP1* or *HNF4A* recapitulated most of the transcription driver changes observed in our CRISPR screens.

**Figure 2.**
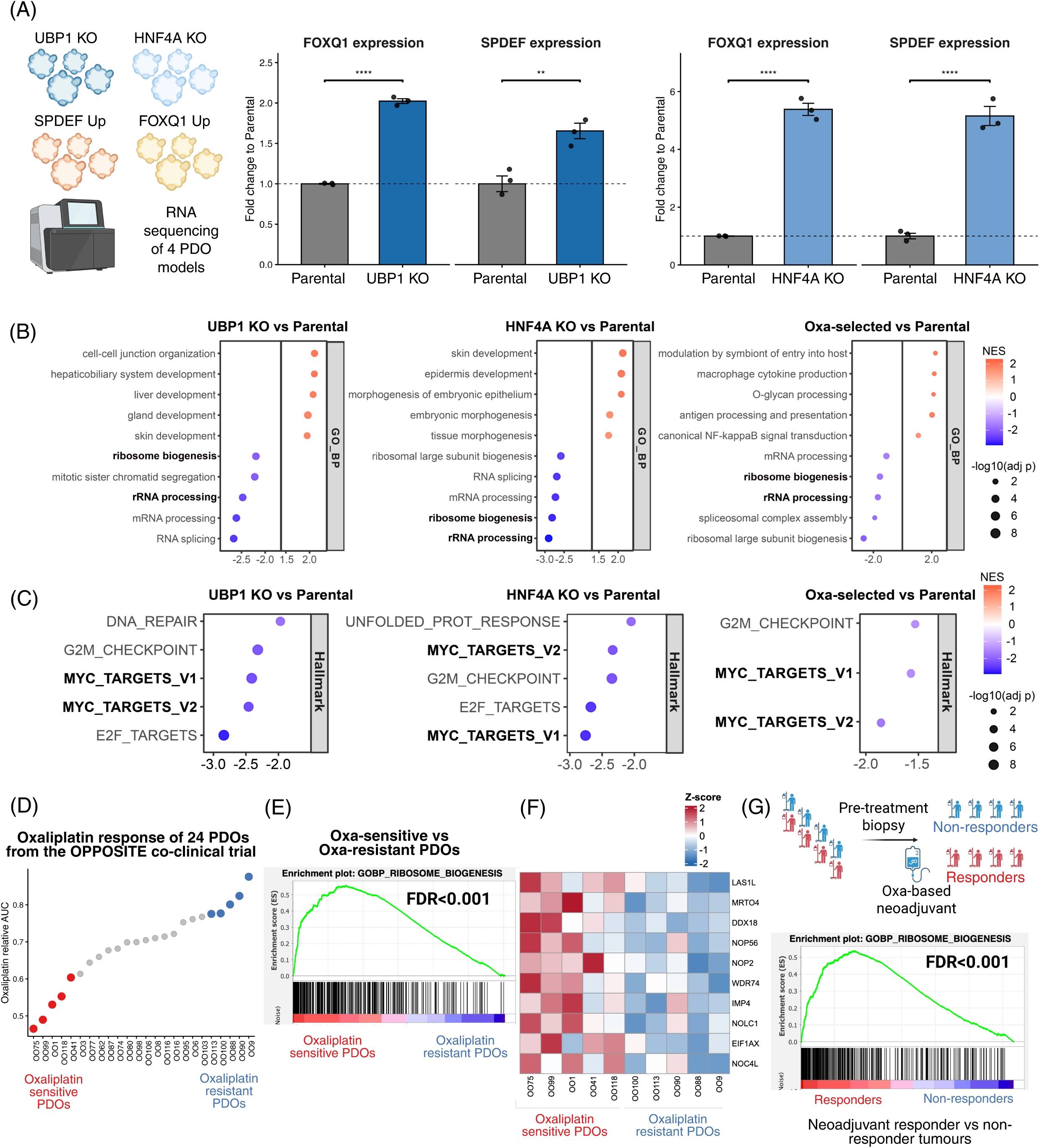
The screen hits are transcriptionally interconnected and drive downregulation of ribosome biogenesis. **(A)** Left panel: a scheme of transcriptomic analysis of each screen hit CRISPR perturbation in sensitive parental organoids. *SPDEF* and *FOXQ1* expression in UBP1 KO (middle panel) and HNF4A KO (right panel) PDO lines. Statistical difference was calculated by Deseq’s Wald t-test (**FDR < 0.01, ****FDR < 0.0001). **(B)** Gene set enrichment analysis (GSEA) of biological processes in UBP1 KO (left panel), HNF4A KO (middle panel), and oxa-selected (right panel) organoid lines compared to parental organoids. Commonly downregulated pathways are highlighted. NES – normalized enrichment score. **(C)** GSEA of hallmark pathways in UBP1 KO (left panel), HNF4A KO (middle panel), and oxa-selected (right panel) organoid lines compared to parental organoids. Commonly downregulated pathways are highlighted. (**D**) Areas under curve (AUC) of 24 PDO lines from the OPPOSITE co-clinical trial upon oxaliplatin treatment. The five most sensitive and the five most resistant PDOs are highlighted. **(E)** GSEA RIBOSOME_BIOGENSIS signature from RNA expression comparing the five most oxaliplatin sensitive with the five most oxaliplatin resistant PDOs from the OPPOSITE trial. **(F)** Heatmap of statistically significant (p < 0.05) MYC-driven ribosome biogenesis factors between oxaliplatin-sensitive and oxaliplatin-resistant PDOs from **(D)**. **(G)** GSEA RIBOSOME_BIOGENSIS signature from RNA expression comparing responder and non-responder tumours to oxaliplatin-based neoadjuvant chemotherapy. FDR – false discovery rate.

To uncover the pathways driven by loss of *UBP1* or *HNF4A*, we analyzed transcriptomic profiles and performed gene set enrichment analysis (GSEA) in the two lines, and compared these genetic models of resistance with the *in vitro-*developed, oxa-selected PDO model. While upregulated biological pathways varied considerably across the three examined datasets, there was significant overlap in downregulated pathways. The ribosome biogenesis and ribosomal RNA (rRNA) processing signatures were downregulated in all three PDOs (Figure 2B). In addition, the hallmarks analysis revealed significant downregulation of MYC target genes – both of the annotated signatures – across all three PDOs (Figure 2C). Since MYC transcription is known to induce ribosome biogenesis^14^, our pathway enrichment analysis reveals downregulation of ribosome biogenesis as a critical common node across all PDO models of oxaliplatin resistance.

Recently, there has been a shift in the conceptualization of oxaliplatin’s mode of action. Several studies suggest that the drug exerts stress on the production and/or function of the translation machinery^8–10^. However, all these data come from 2D cancer cell lines; thus, the clinical relevance of ribosome biogenesis downregulation remains unclear. To examine if such downregulation correlates with resistance to therapy, we analyzed two complementary clinical datasets. First, we recently conducted a co-clinical trial, the OPPOSITE study^15^, which revealed GC PDOs as accurate predictors of chemotherapy response. We profiled the entire cohort of 24 PDOs for oxaliplatin response, then selected the five most sensitive and five most resistant PDOs for further testing (Figure 2D). By comparing transcriptome profiles of the two groups, we found that both ribosome biogenesis and MYC-target genes were significantly downregulated in oxaliplatin-resistant PDOs (Figure 2E, S4A-B). To investigate whether the two identified phenotypes are linked, we curated a set of 38 genes common to both phenotypes. Examining the expression of these genes – that we refer to as “MYC-driven ribosome biogenesis factors” – revealed them to follow a consistent pattern of downregulation in the five resistant organoid lines (Figure 2F). Therefore, ribosome biogenesis, as a determinant of oxaliplatin sensitivity, is explicitly mediated by a MYC-driven mechanism.

Lastly, we interrogated an RNA-seq dataset of GC patients that included tumor samples from responders and non-responders to oxaliplatin-based therapy^16^. While *MYC* amplification has already been shown to be enriched in patients who responded to chemotherapy, we found that the ribosome biogenesis signature was also significantly enriched in responders compared to the non-responders (Figure 2G). In addition, examining pre- and post-treatment data showed that the expression of our curated “MYC-driven ribosome biogenesis factors” was downregulated in the vast majority of non-responder patients after oxaliplatin-based therapy (Figure S5). Altogether, these analyses reveal that all oxaliplatin resistance drivers identified by our CRISPR screens intersect in downregulating MYC-driven ribosome biogenesis. Furthermore, we show that such a shutdown contributes to oxaliplatin resistance/non-response outcome both in a variety of PDOs and in gastric cancer patients.

### Loss of *UBP1* is a common event in gastrointestinal cancers, correlating with poor prognosis

Since the loss-of-function of either UBP1 or HNF4A transcriptionally recapitulated most of the oxaliplatin-resistance changes observed in the CRISPR screens, we investigated the clinical relevance of the loss of either factor. Priority was given to the Cancer Genome Atlas^17–19^ (TCGA) cohorts of gastrointestinal (GI) cancers in which oxaliplatin is currently part of standard therapeutic regimens. While loss of *HNF4A* was an infrequent event across different GI entities (Figure 3A), we found a loss of at least one *UBP1* copy to be remarkably prevalent: 29% of stomach adenocarcinoma (44% in the CIN subtype), 62% of esophageal adenocarcinoma, and 14% of colorectal adenocarcinoma patients harbored the deletion (Figure 3B). These results are in line with the deterministic loss of the 3p arm of chromosome 3 (harboring *UBP1*) early in gastric cancer evolution following *TP53* loss^20^. Given its high mutational prevalence across different GI entities, we focused on the role of UBP1 (Upstream Binding Protein 1) in driving oxaliplatin resistance.

**Figure 3.**
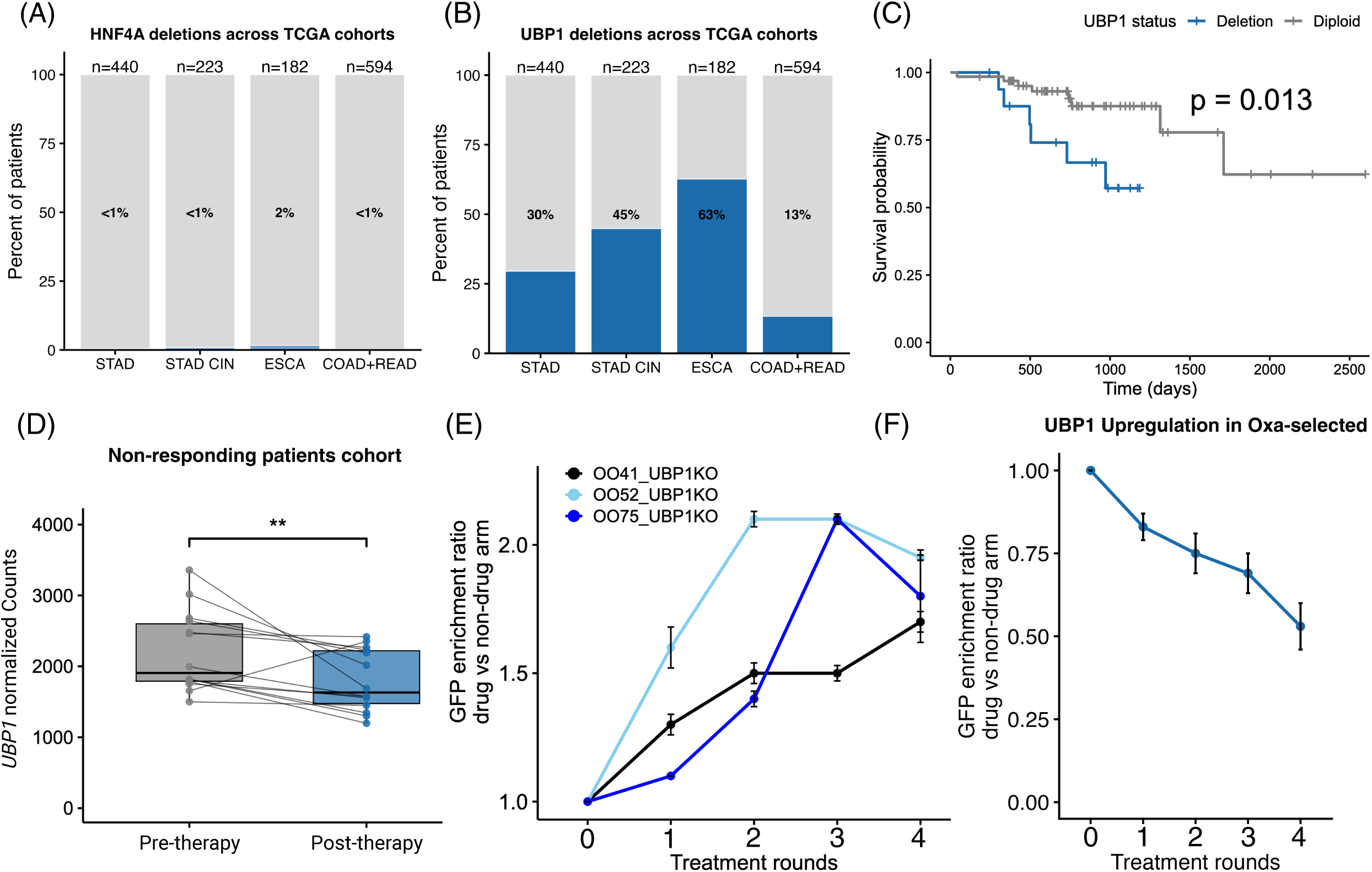
Loss of *UBP1* is a common event in gastrointestinal cancers, correlating with poor prognosis. The frequency of *HNF4A* (**A**) and *UBP1* (**B**) copy-number alterations in the TCGA cohorts: STAD, ESAC, STAD-CIN, and COAD+READ. Deletion designates one or more copy-number loss. **(C)** Kaplan-Meier overall survival of TCGA colorectal (COAD+READ) patients treated with oxaliplatin-based chemotherapy, stratified by *UBP1* deletion versus diploid status; p-value is given by the log-rank test. **(D)** Paired analysis of UBP1 expression in endoscopic biopsies from non-responder patients before (pre-therapy) and after (post-therapy) neoadjuvant oxaliplatin-based chemotherapy. Each line connects one patient (n = 14 pairs); (**p < 0.01) by paired Deseq’s Wald t-test**. (E)** GFP-competition assays in three independent gastric PDOs from different molecular subtypes (OO41, OO52, and OO75 belong to CIN, GS, and CIN subtypes, respectively). The percentage of GFP+ organoids (UBP1 KO) in the drug arm (n=3) is normalized to the non-drug arm. **CIN** Chromosomally unstable; **GS** Genome stable. (**F**) GFP-competition assay of *UBP1*-upregulated (GFP+) organoids versus unmodified parental organoids. The percentage of GFP+ organoids in the drug arm (n=3) is normalized to the non-drug arm. **STAD**: Stomach Adenocarcinoma; **ESAC**: Esophageal Adenocarcinoma; **STAD-CIN**: Chromosomally instable subtype of the Stomach Adenocarcinoma; **COAD+READ**: Combined cohort of Colon Adenocarcinoma (COAD) and Rectum Adenocarcinoma (READ), representing colorectal cancer.

To investigate the impact of *UBP1* deletions on treatment outcomes, we analyzed gastrointestinal cancer patients who have been annotated as having received oxaliplatin. We found that patients with *UBP1* deletion had poorer survival outcomes (log-rank p value 0.013) upon oxaliplatin chemotherapy than those who remained *UBP1* diploid (Figure 3C).

We also interrogated RNA-seq data of pre- and post-treatment tumors in a non-responder cohort of GC patients undergoing oxaliplatin-based therapy^16^. Notably, we found that *UBP1* was downregulated upon treatment in 11 of 14 patients, consistent with its role in driving therapy resistance (Figure 3D). Together, these results demonstrate that *UBP1* loss is a frequent alteration across different GI cancer entities that correlates with poor survival, and that such loss is selected for during therapy.

To test whether loss of UBP1 causes, rather than merely correlates with, oxaliplatin resistance, we aimed to validate its role functionally. First, UBP1 was the most robust screen hit, with four out of five of its guides scoring among the top 1% of all gRNAs in the CRISPR-Cas9 screen (Figure S6A). Second, we tested UBP1 resistance-driving capacity utilizing our organoid biobank. Strikingly, UBP1 KO caused oxaliplatin resistance in all three examined PDOs, two of which belonged to the chromosomally unstable subtype (CIN: OO41 and OO75), while one belonged to the genome-stable subtype (GS: OO52; Figure 3E). Of note, UBP1 KO did not cause an increase in proliferation and/or survival in the absence of oxaliplatin in any of the tested organoids (Figure S6B). These results demonstrate that the genetic loss of *UBP1* commonly causes oxaliplatin resistance, a broader mechanism not confined to the PDO line used for the screening.

Finally, we examined whether restoring *UBP1* expression in oxa-selected DD109 would resensitize them to oxaliplatin. Namely, *UBP1* was included in the CRISPR-Cas9 screen due to a 3p chromosome deletion in the resistant model (Figure S7A-B). We first confirmed that the oxa-selected line showed a ∼3-fold decrease in *UBP1* expression compared to parental DD109 (Figure S7C). Subsequently, using the CRISPRa system, we selected a gRNA that upregulated *UBP1* expression in oxa-selected DD109 organoids, resulting in expression levels comparable to those in parental organoids (Figure S7D). Using the same GFP competition assay described earlier, we confirmed potent resensitization of previously resistant organoids upon upregulation of *UBP1* expression (Figure 3F). In summary, loss of UBP1 not only correlates with oxaliplatin resistance in large patient cohorts but also causes such resistance across a wide range of GC PDOs, while restoring its expression induces oxaliplatin sensitivity.

### *UBP1* loss causes a diminished *MAX* expression and protects from oxaliplatin-induced nucleolar damage

The extensive transcriptomics analysis of PDOs from the co-clinical trial revealed that MYC-driven ribosome biogenesis was downregulated in the resistant PDOs. To assess whether loss of UBP1 caused downregulation of the same signature, we analyzed transcriptomes of UBP1 KO organoids and the *in vitro-*developed oxa-selected model, in which one copy of the *UBP1* gene has been lost. Indeed, more than 70% of “MYC-driven ribosome biogenesis factors” were significantly downregulated in both resistant models compared to the parental line (FDR < 0.05, Figure 4A).

**Figure 4.**
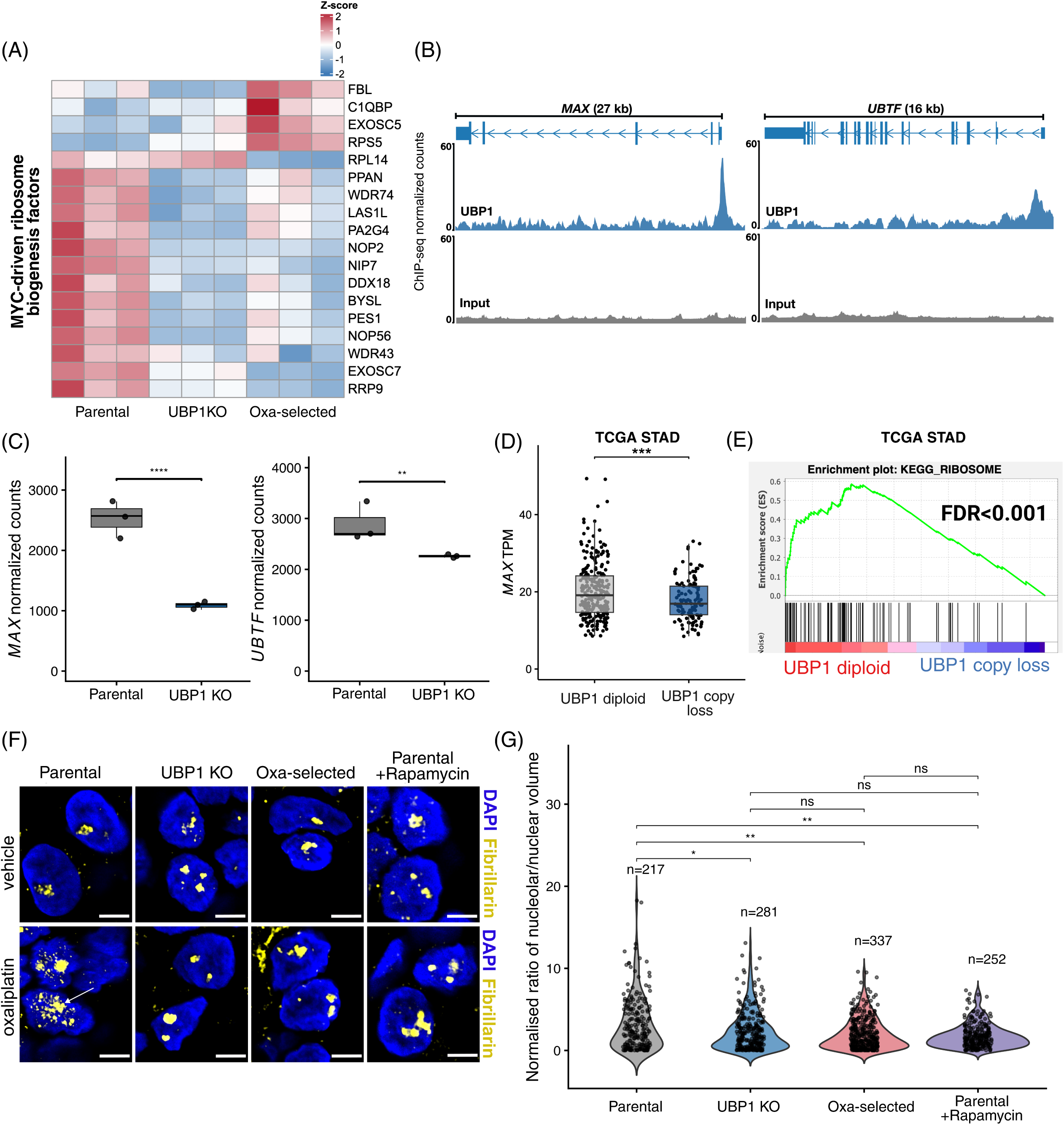
*UBP1* loss causes a diminished *MAX* expression and protects against oxaliplatin-induced nucleolar damage. **(A)** Heatmap of statistically significant genes (FDR < 0.05) between parental and oxa-selected PDOs showing RNA expression for a curated set of “MYC-driven ribosome biogenesis factors” in parental, UBP1 KO, and oxa-selected organoid lines. **(B)** CHIP-seq reads of UBP1 binding to the promoter region of *MAX* (left panel) and *UBTF* (right panel). **(C)** RNA-seq quantification of *MAX* (left panel) and *UBTF* (right panel) expression with adjusted p-values from DESeq Wald test shown for parental vs. UBP1 KO organoids (**FDR < 0.01 and ****FDR < 0.0001). **(D)** Comparison of RNA-seq quantification of *MAX* in *UBP1* diploid cohort and *UBP1* copy loss cohort in TCGA STAD patients. Statistical difference was calculated by a two-sided t-test (***p < 0.001). **STAD**: Stomach Adenocarcinoma. **(E)** GSEA RIBOSOME signature between *UBP1* diploid and *UBP1* copy loss TCGA STAD patient cohorts. FDR – false discovery rate. (**F**) Immunofluorescence (IF) images depict nucleoli (Fibrillarin) and nuclei (DAPI) of parental (first column), UBP1 KO (second column), oxa-selected (third column), and parental organoids treated with rapamycin (20 μM, fourth column) in the presence (bottom panels) and absence (upper panels) of oxaliplatin (4 μM). Arrow points to a loss of nucleolar integrity in parental organoids treated with oxaliplatin. Scale bars depict 5 μm. (**G**) Quantification of nucleolar volume changes in PDO lines shown in (**F**). The share of nucleolar to nuclear volume upon oxaliplatin treatment has been normalized for each PDO line to the control treatment. Statistical comparisons between different lines were performed by the Wilcoxon test (*p < 0.05 and **p < 0.01).

We next sought to elucidate how *UBP1* loss causes diminished MYC-driven ribosome biogenesis. Peak annotation of UBP1 binding sites in the ChIP-seq experiment revealed UBP1 enrichment around promoter regions and transcription start sites, in agreement with its transcription factor activity (Figure S8A-B, Supplementary Table S8). Upon a closer examination of key genes involved in driving ribosome biogenesis transcription – *UBTF, MYC, and MAX*^25^ – we found that UBP1 strongly bound the promoter region of *MAX and* moderately bound the *UBTF* promoter (Figure 4B). *MAX* is a co-transcriptional factor required for MYC target binding^26^, whereas UBTF is the main organizer of rDNA open conformation and an indispensable factor in rRNA transcription^27^. While both genes were downregulated in UBP1 KO organoids, their expression correlated with UBP1 binding strength, with *MAX* downregulation being more pronounced than *UBTF* downregulation (Figure 4C). Furthermore, in the oxa-selected resistant model, a partial loss of *UBP1* was followed by downregulation of *MAX* expression, but not of *UBTF* expression (Figure S9A-B). Therefore, our data suggest that UBP1 is a critical driver of the master regulators of ribosome biogenesis, and in particular of *MAX*.

To assess the robustness of these findings, we examined the TCGA stomach adenocarcinoma (STAD) cohort and divided patients into those who lost at least one *UBP1* copy and those who remained diploid for the gene. We first confirmed that loss of one copy of *UBP1* significantly diminished its expression (Figure S10). More importantly, loss of *UBP1* was associated with significant downregulation of *MAX* compared with patients with diploid copy number (Figure 4D). Furthermore, when we compared the transcriptomes of two patient cohorts, the RIBOSOME signature was significantly downregulated in patients haploid for *UBP1* (Figure 4E). Taken together, these data indicate that a loss of *UBP1* leads to downregulation of ribosome biogenesis through direct regulation of *MAX* expression and, thereby, MYC activity.

Lastly, we sought to link ribosome biogenesis and nucleolar stress induced by oxaliplatin. We examined nucleolar dynamics by tracing fibrillarin, a constitutive nucleolar structural protein^28^, in different resistance models in the presence and absence of oxaliplatin. In parental sensitive organoids, treatment with oxaliplatin resulted in major disruption of nucleolar structure, leading to increased nucleolar volume and loss of nucleolar integrity (Figure 4F-G). In UBP1 KO and oxa-selected resistant models, such disruption by oxaliplatin was significantly lower or absent (Figure 4F-G), suggesting that downregulation of ribosome biogenesis protects from oxaliplatin-induced nucleolar stress. To directly test for this hypothesis, we co-treated parental organoids with oxaliplatin and rapamycin, a known blocker of mTOR and ribosomal biogenesis^29,30^. Rapamycin protected organoids from nucleolar disassembly (Figure 4F), and quantification of immunofluorescence imaging revealed that UBP1 KO and rapamycin displayed similar protective effects (Figure 4G). Taken together, our results indicate that loss of *UBP1* leads to downregulation of MYC-driven ribosome biogenesis, and thereby, protects cells from oxaliplatin-induced nucleolar stress.

### Activation of translation initiation, mediated by 4EBP1 loss-of-function, compensates for ribosome biogenesis downregulation

Our discovery that UBP1-driven, and consequently, MYC-driven ribosome biogenesis downregulation is associated with oxaliplatin resistance cannot be easily reconciled with a cancer cell’s high demand or even addiction to protein synthesis. To explore this paradox, we examined the translation dynamics in oxaliplatin resistance models. First, we found that the key translation regulator 4EBP1, a well-described translation initiation repressor^31^, was downregulated in UBP1 KO organoids, as well as in the oxa-selected model (which carries a *UBP1* deletion) compared to parental organoids, on the RNA (Figure 5A) and protein levels (Figure 5B). 4EBP1 is a central node in translation regulation across many pathways, and especially the mTOR pathway, where mTORC1 regulates 4EBP1 activity through inhibitory phosphorylation^30^. Indeed, we observed an increase in phosphorylation status of 4EBP1 across the resistance models, concomitant with its downregulation (Figure 5B). Both of these events – total protein downregulation and functional inactivation – lead to the release of the eIF4E initiation factor from 4EBP1 and, consequently, the activation of translation^30^.

**Figure 5.**
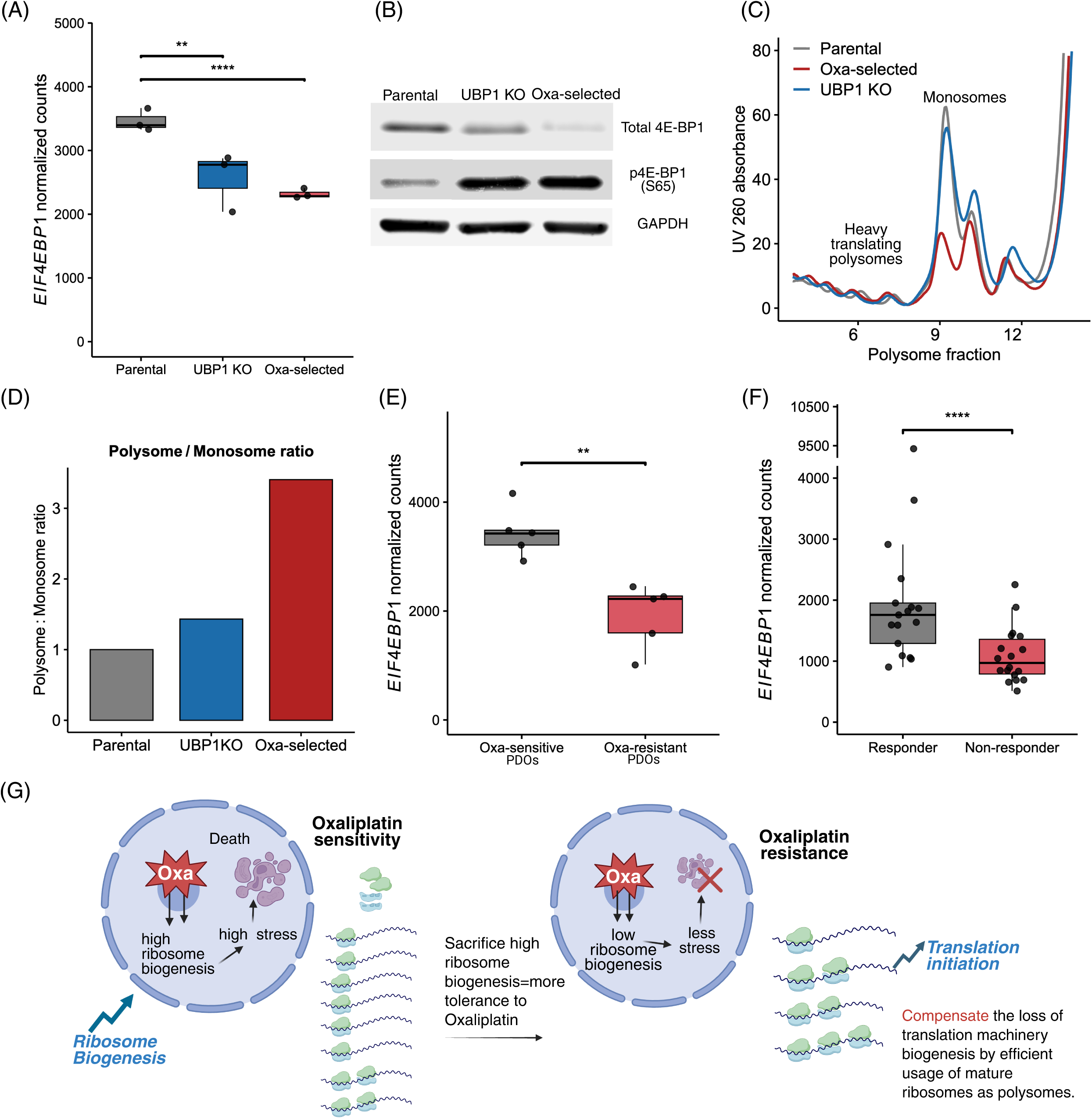
Activation of translation initiation by 4EBP1 loss-of-function compensates for downregulation of ribosome biogenesis. **(A)** RNA-seq quantification of *EIF4EBP1* expression with adjusted p-values from Deseq’s Wald test shown for parental vs. UBP1 KO and parental vs. oxa-selected organoid lines (****FDR < 0.0001, **FDR < 0.01). **(B)** Western blot analysis of total and phospho-4EBP1 (Ser65) in parental, UBP1 KO, and oxa-selected PDOs. **(C)** Representative polysome profiles (A□□□) from DD109 PDOs: parental, UBP1 KO, and oxa-selected. Monosome and heavy polysome peaks are annotated. **(D)** Quantification of the polysome-to-monosome ratio (area under the curve) in organoid resistance models. **(E)** RNA-seq quantification of *EIF4EBP1* expression with p-values from Deseq’s Wald test between the five most sensitive and the five most resistant organoids from the OPPOSITE study (**p < 0.01). **(F)** RNA-seq quantification of *EIF4EBP1* expression with p-values from Deseq’s Wald test between responder and non-responder tumour biopsies before oxaliplatin-based neoadjuvant chemotherapy stratified by clinical response (****p < 0.0001). **(G)** Proposed model: crucial steps are highlighted.

We next aimed to functionally estimate the consequences of translation initiation activation by conducting a polysome profiling experiment^32^ in different organoid lines. Comparing the profiles of the parental, UBP1 KO, and oxa-selected models (Figure 5C and Supplementary File 1) revealed an increase in the polysome-to-monosome ratio in oxaliplatin-resistant models (Figure 5D), indicating that a more efficient use of ribosomes is a feature of resistant organoids.

Finally, to investigate the clinical relevance of 4EBP1 loss-of-function, we turned to the PDO biobank from the OPPOSITE trial. Comparing transcriptomes of the five most resistant PDOs to the five most sensitive to oxaliplatin, we observed a significant downregulation of *EIF4EBP1* (coding for 4EBP1) in resistant organoids (Figure 5E). This finding was further supported by comparing RNA-seq data from pre-treatment biopsies between responder and non-responder tumors^16^, in which the non-responder cohort exhibited lower *EIF4EBP1* expression (Figure 5F). Compensatory reliance on translation initiation, therefore, seems to be a common route of resistance in gastric cancer.

In sum, we propose a working model in which oxaliplatin sensitivity of cancer cells is primarily mediated by hyperactive MYC-driven ribosome biogenesis, which, upon oxaliplatin treatment, leads to a high level of ribosome biogenesis/nucleolar stress, resulting in cell death. In oxaliplatin-resistant tumors, conversely, tumor cells sacrifice a high level of ribosome biogenesis, becoming more tolerant to the drug. Simultaneously, the tumors resort to heightened translation initiation dynamics to compensate for the loss of translation machinery biogenesis, thereby maintaining protein synthesis levels required for proliferation and growth. A graphical representation of the model is presented in Figure 5G.

### Pharmacological restoration of 4EBP1 function via 4EGI resensitizes resistant organoid models

Our findings prompted us to consider whether restoring 4EBP1 function could reverse oxaliplatin resistance. To investigate this hypothesis, we used 4EGI, a pharmacological mimic of 4EBP1 that binds to the rate-limiting translation initiation factor eIF4E^33–35^. Simultaneously, the compound enhances the binding of endogenous 4EBP1 to eIF4E, thereby inhibiting translation initiation and global translation^33^. To assess the efficacy of combining 4EGI with oxaliplatin, we used an organoid formation assay, as well as a cell death assay (Figure S11A-B).

We have shown that *UBP1* knockout protected against oxaliplatin-induced nucleolar stress (Figure 4F-G), but simultaneously rendered cells more reliant on translation initiation (Figure 5B-C). Indeed, the colony formation assay confirmed that UBP1 KO organoids were less affected by oxaliplatin compared to the parental line, while the opposite was true in the case of 4EGI treatment (Figure 6A, left and middle panels). Thus, 4EGI targeting reveals a specific vulnerability in organoids that compensate for ribosome biogenesis downregulation by upregulating translation initiation. Furthermore, the combination of oxaliplatin and 4EGI decreased organoid-forming capacity compared with either single-agent treatment in both the parental DD109 and the two resistance models (Figure 6A). However, the quantification of excess death using the Bliss independence model revealed a merely additive effect in the parental line, whereas in both resistant PDO models a synergistic effect was observed (Figure 6B). A similar synergism was also revealed in the most oxaliplatin-resistant PDO line from the OPPOSITE trial (Figure S12A-B). These findings validate our hypothesis that restoring 4EBP1 functionality and inhibiting translation with 4EGI is an effective strategy for resensitizing otherwise resistant models.

**Figure 6.**
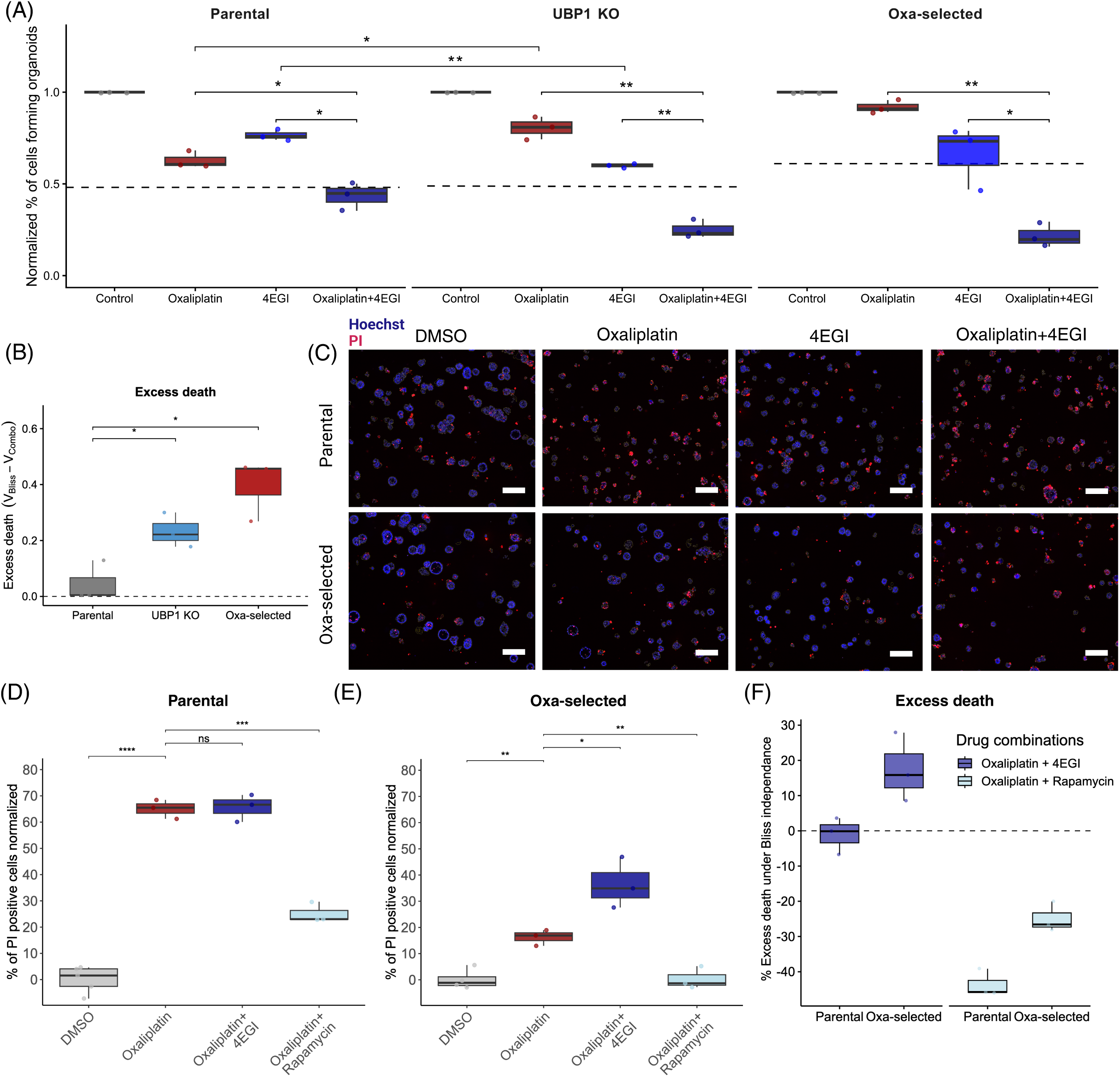
Pharmacological mimicry of 4EBP1 via 4EGI reverses resistance in PDO models. **(A)** Organoid formation assays in the parental DD109 (left panel), UBP1 KO (middle panel), and oxa-selected (right panel) PDOs treated with control (DMSO), oxaliplatin, 4EGI, or a combination of both (n=3). Organoid-forming efficiency is normalized to the DMSO vehicle. The dashed line represents the expected additive effect under the Bliss independence model. Horizontal bars denote paired (within the same PDO) and unpaired (between two different PDOs) two-sided Student’s t-test comparisons (*p < 0.05, **p < 0.01, ***p < 0.001). **(B)** Excess death is calculated as the difference between expected and observed organoid formation capacity in the parental and the two resistant models under the Bliss independence model. Horizontal bars denote two-sided Student’s t-test comparisons (*p < 0.05). **(C)** Representative images of cytotoxicity assay by propidium iodide (PI, red) uptake in parental PDO line (upper panels, n=3) or oxa-selected PDO (bottom panels, n=3) following indicated treatments. Cell nuclei are marked by Hoechst (blue). Scale bars represent 200 μm. **(D, E)** Quantification of apoptosis by PI uptake in parental **(D)** or oxa-selected **(E)** PDOs following indicated treatments. Data are expressed as a percentage of PI-positive cells, normalized to DMSO (vehicle). Concentrations used were oxaliplatin (10 μM), 4EGI (50 μM), and rapamycin (20 μM). Statistical significance was determined by two-sided Student’s t-test: *p < 0.05, **p < 0.01, ***p < 0.001, ****p < 0.0001; “ns” = not significant. **(F)** Excess cell death under the Bliss independence for two oxaliplatin-based drug combinations: oxaliplatin + 4EGI, and oxaliplatin + rapamycin. Bars represent median with interquartile range (n = 3).

Oxaliplatin has been variably described as either a ribosome biogenesis stressor or a general translation stressor^9,36^. However, our data indicate that while high ribosome biogenesis sensitizes tumors to oxaliplatin, high translation rates are associated with resistance. Therefore, we sought to investigate the specific contributions of two major translation arms – ribosome biogenesis and translation initiation – to oxaliplatin-induced cell death. To do so, we assessed the efficacy of oxaliplatin/4EGI combination compared to oxaliplatin combination with rapamycin, an inhibitor of both ribosome biogenesis and translation initiation^29,30^, for which we have already shown protective effects from oxaliplatin-induced nucleolar stress (Figure 4F-G). We employed an image-based assay, and the detailed readout allowed us to assess both cell death (PI signal) and organoid growth arrest (number of segmented nuclei, Figures S13 and S14). While 4EGI and rapamycin alone did not cause significant levels of death compared to oxaliplatin, they caused a comparable, yet prominent inhibition of growth in both the parental and oxa-selected organoids (Figure S13A-B), indicating that the selected doses were functionally relevant.

The cell death assay further replicated previously observed additive effects of oxaliplatin/4EGI combination in parental PDO, as well as the synergistic effects in the oxa-selected resistant line (Figure 6C-F, Figure S14A-B). However, rapamycin displayed a clear and strong protective effect in both assessed lines, reducing the number of PI-positive organoids when combined with oxaliplatin compared with oxaliplatin alone (Figure 6D-E and S15). The excess death analysis further revealed a large extent of rescue provided by rapamycin (Figure 6F), in agreement with the drug’s protection of nucleolar integrity upon oxaliplatin treatment. Together, these findings reveal that: 1) 4EGI shows a unique and specific synergy with oxaliplatin only in the context of translation initiation addiction found in oxaliplatin-resistant organoids, and 2) that a successful resensitization requires selective inhibition of translation initiation only, since any inhibition of ribosome biogenesis will instead protect against oxaliplatin-induced nucleolar stress, and consequently against oxaliplatin’s toxicity.

### Oxaliplatin resistance acquired *in vivo* mirrors the key molecular features observed *in vitro* and exhibits vulnerability to 4EGI

The translational potential of 4EGI prompted us to consider whether our findings could be replicated in a model in which oxaliplatin resistance emerged *in vivo*. We therefore obtained a pair of PDOs, generated from the same patient before (OO99) and after four cycles of FLOT treatment (OO99cT). The pre-treatment line OO99 was among the oxaliplatin-sensitive PDOs in the OPPOSITE study^15^; however, we observed a decrease in oxaliplatin sensitivity in the post-treatment OO99cT line (Figure 7A). This isogenic pair, then, reflects an *in vivo* evolution of oxaliplatin resistance following the standard-of-care treatment (Figure 7A).

**Figure 7.**
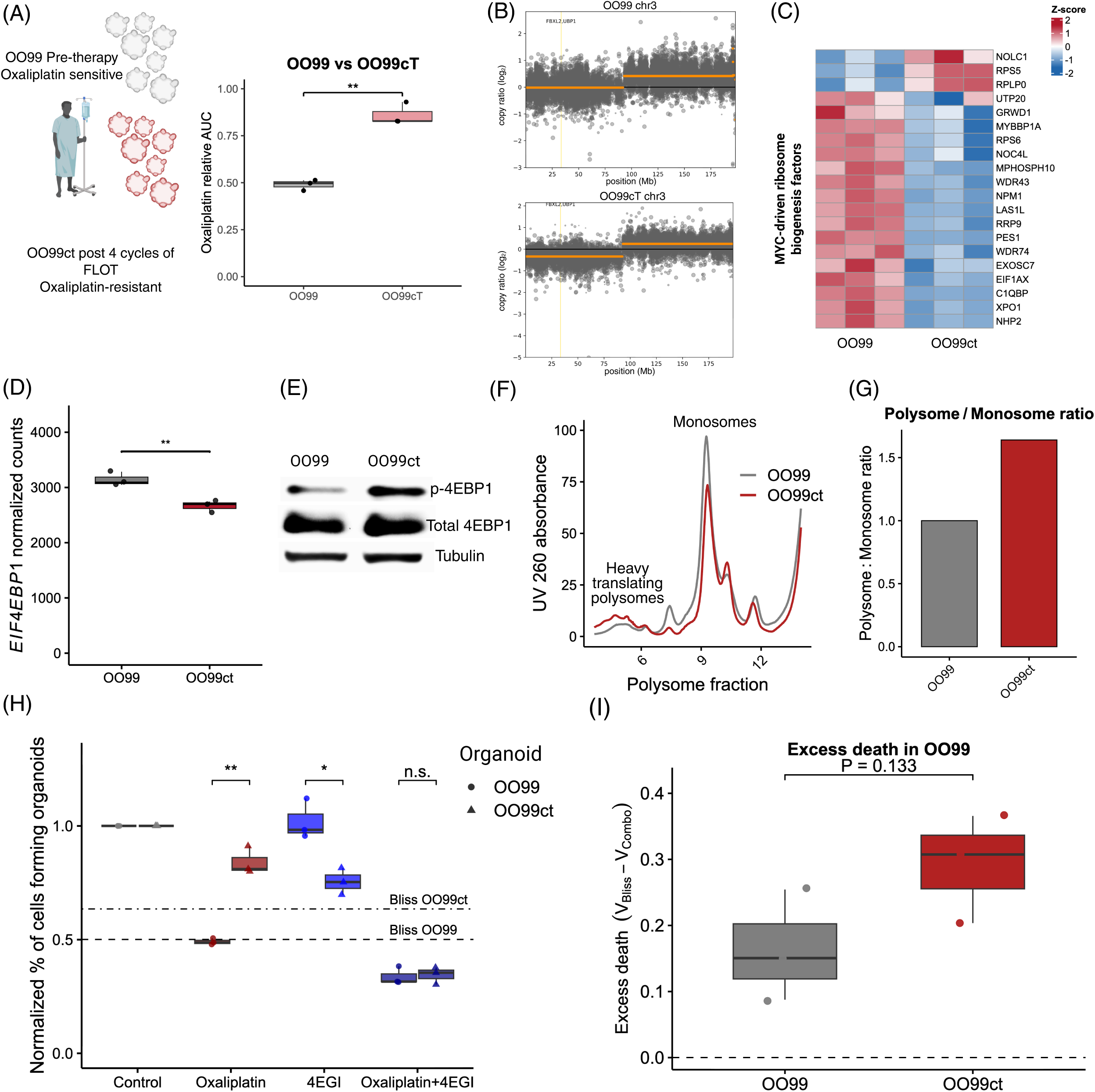
Oxaliplatin resistance acquired *in vivo* mirrors the key molecular features observed *in vitro* and exhibits marked sensitivity to 4EGI. **(A)** A schematic of pretreatment biopsy-derived OO99 organoids and from the same patient after four cycles of FLOT chemotherapy (OO99ct, left); The difference in oxaliplatin sensitivity between the two PDOs by relative AUC (n=3; right). **(B)** Chromosome-wide copy-number profiles (log□ copy ratio) across chromosome 3 for OO99 and OO99ct organoids, with the *UBP1* locus highlighted. **(C)** Heatmap of Z-scored RNA expression for a curated MYC-driven ribosome biogenesis signature in OO99 versus OO99ct organoids. Statistically significant genes (FDR < 0.05) are plotted. **(D)** RNA-seq quantification of *EIF4EBP1* expression in OO99 and OO99ct organoids with adjusted Deseq’s Wald test p-value indicated above the comparison (**FDR < 0.01). **(E)** Western blot analysis of total and phospho-4EBP1 (Ser65) in OO99 and OO99ct organoids. **(F)** Polysome profiling from OO99 and OO99ct organoids, with monosome and heavy-polysome peaks annotated. **(G)** Quantification of the polysome-to-monosome ratio as AUC, normalized to OO99. **(H)** Organoid formation assays in OO99 and OO99ct. Organoids were treated with DMSO control, oxaliplatin (6.5 μM), 4EGI (33 μM), or the combination. Organoid-forming efficiency is normalized to control (dashed line, n = 3); statistical comparisons were performed by two-sided Student’s t-test. Dashed lines represent additive effects under the Bliss independence model. **(I)** Comparison of excess death under the Bliss independence model in oxaliplatin/4EGI combination between OO99 and OO99ct organoids (n = 3). *p < 0.05, **p < 0.01, ***p < 0.001, “n.s.” - not significant.

Intriguingly, we found a chromosome 3p deletion enriched in the post-therapy PDO line, similar to the *in vitro-*selected resistance model (Figure 7B). In addition, RNA-seq analysis showed a downregulation of the majority of genes of the curated “MYC-driven ribosome biogenesis factors” set in the post-treatment PDO (Figure 7C). To evaluate the translational relevance of these findings, we first examined *EIF4EBP1* expression. We found it to be transcriptionally downregulated in the post-therapy PDO, as in the various *in vitro* models (Figure 7D). Furthermore, western blot analysis revealed that phosphorylated (inhibited) 4EBP1 was more abundant in OO99cT, consistent with previous findings (Figure 7E). Consequently, the polysome profiling of pre- and post-therapy PDOs showed that the polysome-to-monosome ratio was higher in OO99ct organoids (Figure 7F-G and Supplementary file S2). This suggests that *in vivo* resistance development also led to increased translation initiation, which was mediated by the loss of 4EBP1 function.

Finally, we sought to evaluate the efficacy of the oxaliplatin/4EGI combination in the isogenic organoid pair. Strikingly, the evolution of resistance to oxaliplatin was followed by an increase in 4EGI sensitivity, a feature of reliance on translation initiation in the absence of high ribosome biogenesis (Figure 7H). Of note, the post-treatment PDO line, which had shown 30% less cell death under oxaliplatin alone than the pre-treatment PDO, became equally responsive when treated with oxaliplatin/4EGI combination (Figure 7H). Furthermore, while the combinatorial treatment exhibited a synergistic effect in both pre- and post-treatment PDOs (Figure 7H), the excess death under the Bliss independence model was higher in the post-treatment sample, although the effect was not significant (Figure 7I). Taken together, these results provide evidence that the proposed therapeutic strategy demonstrates efficacy also in tumor cells that have evolved resistance *in vivo* under clinically relevant treatment conditions.

## Discussion

In this study, we provide key insights into the determinants of sensitivity, resistance, and resensitization to one of the most frequently used chemotherapeutic agents in oncology — oxaliplatin — paving the way for better treatment strategies. Our work builds on an emerging paradigm positing oxaliplatin as a ribosome biogenesis and/or translation stressor, adding a critical distinction. Employing a multi-platform approach comprising patient-derived organoids, CRISPR screening, and a wide range of multi-omics tumor datasets, we analyzed the molecular mechanisms governing oxaliplatin response and resistance in gastric cancer. Crucially, we identify a common genetic marker of resistance and uncover a targetable molecular vulnerability in resistant tumor cells.

One of the identified screen hits was shown to be commonly deleted in GC patients. We discovered that loss of *UBP1* is a major determinant of oxaliplatin resistance, characterized by a diminished *MYC*-driven ribosome biogenesis. While our work suggests that *UBP1* loss reduces MYC-driven transcription via the MYC cofactor MAX, it is worth noting that there is already evidence linking MYC and UBP1 directly. Namely, *UBP1* was first described more than 30 years ago as *LBP1*, a crucial repressor of HIV transcription^37^. Later, it was found that MYC, together with UBP1, induced synergistic repression of HIV^38^. Therefore, it remains to be seen whether MYC and UBP1 interact directly to activate the ribosome biogenesis program.

Interestingly, a recent study characterizing the deterministic evolution of GC PDOs following *TP53* loss identified chromosome 3p deletion, harboring *UBP1*, as one of the earliest and most recurrent events in gastric tumorigenesis^20^. In line with these findings, this deletion was highly prevalent in the gastrointestinal cancer cohorts we analyzed, reaching up to 62% of patients in esophageal adenocarcinoma. The *FHIT* gene was proposed as the driving force behind the evolutionary advantage of tumor clones with 3p loss^20^. Our data are in agreement with this, as *UBP1* loss does not confer a selective advantage in the absence of oxaliplatin. In three different PDOs, UBP1 KO resulted in only a mild, yet notable, growth retardation in the absence of the drug (Figure S2E). Thus, it appears that the early 3p loss, favored due to *FHIT*, creates a “bystander” *UBP1* loss, which subsequently becomes a substrate for natural selection during treatment.

Our comprehensive CRISPR screens identified four transcription factors driving the resistance to oxaliplatin. Remarkably, all of the factors were transcriptionally interconnected, and the common pathway downregulated across all resistance models was ribosome biogenesis. The initial observation of oxaliplatin’s notable efficacy in cells with high ribosome activity was made in the context of colorectal cancer cell lines^9^. There, the phenomenon was attributed to *APC* mutations, which are known to induce hyperactivation of protein synthesis^39^. While this analysis revealed a hitherto unknown mechanism of oxaliplatin action, it did not discriminate between two critical nodes of protein production: the biogenesis of the translation machinery and the employment of such machinery in the translation of mRNAs. Namely, the authors interchangeably used two terms – “ribosome biogenesis” and “translation addiction” – maintaining that an increase in either correlates with sensitivity to the drug. Our study elucidates a crucial difference. First, we show that *MYC* activity is a key determinant of oxaliplatin therapeutic efficacy in GC. MYC is a known activator of ribosome biogenesis^14^, and *MYC* activation in stomach tissue has been attributed to the unleashing of a Wnt oncogenic program, in striking similarity to APC loss-of-function^40^. However, we provide evidence that translation initiation activity is a compensatory mechanism contributing to the resistance, primarily via the functional loss of 4EBP1. It is worth noting that MYC has been shown to bind the *EIF4EBP1* promoter^41^, providing a possible mechanistic link. Overall, in our rendering, the sacrifice of downregulating ribosome biogenesis is (partially) recuperated by more efficient usage of remaining ribosomes in the resistant cells. The present study, therefore, cautions against using totalizing descriptors such as “translation addiction” and instead proposes two distinct nodes in the process of protein production that play opposing roles in driving resistance to oxaliplatin: ribosome biogenesis and translation initiation.

Importantly, the new conceptualization of oxaliplatin as a ribosome biogenesis stressor does not, in itself, offer a clear perspective on the problem of resistance. Here, by identifying a compensatory mechanism in resistant cells, we provide evidence of cross-addiction from high ribosome biogenesis to high translation initiation, with direct therapeutic implications. We show that, in addition to downregulating ribosome biogenesis, *UBP1* loss also activates translation initiation, rendering resistant cells vulnerable. Specifically, the combined action of 4EGI with oxaliplatin demonstrated remarkable synergism in killing translation initiation-addicted oxaliplatin-resistant PDO models. Notably, rapamycin – a dual blocker of both ribosome biogenesis and translation initiation – exhibited almost a complete rescue of oxaliplatin cytotoxicity. Thus, effective resensitization to oxaliplatin requires selective inhibition of translation initiation, whereas any inhibition of ribosome biogenesis mitigates cytotoxicity by protecting against oxaliplatin-induced nucleolar stress.

Confirming our findings from *in vitro*-developed models in an *in vivo*-developed model of oxaliplatin resistance underscores their generalizability. Notably, the oxaliplatin/4EGI combination was comparably effective across *in vitro* and *in vivo* resistance models, highlighting its translational potential in tumors that developed oxaliplatin resistance under treatment pressure. This is of utmost interest in the setting of gastric cancer, in which patients receive oxaliplatin-based chemotherapy (i.e., FLOT) before and after resection of the primary tumor, giving tumor cells ample time to develop resistance via *UBP1* loss. Consequently, the present work warrants further testing of 4EGI in a clinical trial. In sum, our findings demonstrate that understanding and targeting translation initiation addiction could offer a powerful route toward more effective and tailored treatment regimens for gastric cancer patients.

## Methods

### Patient samples and organoid culture

Primary gastric cancer specimens were obtained from patients undergoing surgery at the University Hospital Carl Gustav Carus, TU Dresden, or University Heidelberg with written informed consent (ethical vote numbers EK451122014, EK169052018, and S-686/2017).

PDOs were generated and cultivated as described previously^11,15,42^ Briefly, endoscopic biopsies were minced, enzymatically digested using dispase II (Roche) and collagenase XI (Sigma-Aldrich), washed three times with antibiotics-containing medium (DMEM/F12, 10 mM HEPES (Thermo Fisher Scientific), 1x GlutaMAX™ (Thermo Fischer Scientific), 100 μg/ml Primocin (Invivogen)) and seeded in Matrigel (Corning). Gastric organoid medium (50% WNT-conditioned medium, 10% R-spondin-conditioned medium, 10% Noggin-conditioned medium, 1x B27 (Invitrogen), 10 mM nicotinamide (Sigma-Aldrich), 1 mM N-acetyl-L-cysteine (Sigma-Aldrich), 200 ng/ml human recombinant fibroblast growth factor hFGF10 (Preprotech), 50 ng/ml recombinant mouse epithelial growth factor mEGF (Invitrogen), 2 μM A83-01 (Tocris Bioscience), 100 μg/ml Primocin (Invivogen), 1 nM human gastrin (Sigma-Aldrich)) was overlaid and supplemented with 10 μM Y-27632 (Sigma-Aldrich) for the first 2-6 passages. Depending on the growth rate, organoids were passaged two or three times a week.

### CRISPR library design

CRISPR-Cas9 knockout (KO) and CRISPR activation (CRISPRa) libraries were designed to identify genes associated with oxaliplatin resistance in the gastric PDO line DD109 and its oxaliplatin-resistant derivative, DD109 oxa-selected.

For the KO screen, candidate genes were prioritized if meeting at least one of the following criteria: (i) strongly downregulated in RNA-seq (fold change > 4, p_adjusted_ <0.0001); (ii) moderately downregulated and correlated with poor survival in TCGA-STAD (p_adjusted_ <0.0001, correlation p < 0.05); (iii) concordant loss in exome copy number analysis; (iv) protein-truncating mutations with significant VAF shifts (> 45% for novel mutations, > 30% for LOH); or (v) recurrent deleterious missense mutations predicted by SIFT^43^ Full list of differentially expressed genes, mutations, as well as copy number alterations between parental and oxa-selected PDOs is provided in Supplementary Tables S1-3, respectively. The annotation of gene category is given in the KO screen results as Alteration category (Supplementary Table S6).

For the CRISPRa screen, candidates included: (i) most upregulated genes (top 100 highest log_2_FC with p_adjusted_<0.001), (ii) most significantly upregulated genes (top 150 lowest p_adjusted_ with fold change more than 1.5) in DD109 oxa-selected organoids, (iii) genes associated with exome copy number gain and upregulation (fold change >1.5 and p_adjusted_ <0.001), (iv) those whose expression correlated with patient survival in TCGA-STAD (fold change >1.5, expression-adjusted p < 0.0001, correlation p < 0.05). Only genes with ≥1 FPKM in resistant PDOs were included to avoid false positives. The annotation of each gene category is provided in CRISPRa screen results (Supplementary Table S7).

gRNAs were designed using the Brunello library and CRISPRPick^44^ selecting five gRNAs per gene. KO gRNAs were prioritized based on on-target score, coding sequence coverage, and proximity to the translation start site, while CRISPRa gRNAs were chosen from the promoter-proximal region (−300 to 0 bp from TSS). Full lists of gRNA sequences are supplied in Supplementary Tables S4-5.

### Cloning and lentiviral production

Custom oligonucleotide pools (Twist Bioscience, Supplementary Table S1-2) encoding gRNAs were PCR-amplified and cloned into lentiviral vectors using Golden Gate assembly. For KO screens, gRNAs were cloned into GFP- or puromycin-selectable vectors, as previously described^12^ (pL.CRISPR-GFP, pL-CRISPR-puro backbones). For CRISPRa screens, DD109 PDO was first engineered to stably express dCas9-VPR, a gift from Kristen Brennand (Addgene plasmid #99373). To enable dual selection, previously described universal plasmids^12^ were used to deliver gRNAs, either in conjunction with GFP or blasticidin. Golden gate cloning reaction was performed using 100 ng pL.CRISPR or universal vector in 1x Tango buffer (Thermo Fisher), 50 nM DTT, 0.5 μL T4 ligase (New England Biolabs), 1 μL Esp3 (Thermo Fisher) and gRNA PCR pool in 3:1 molar ratio to the CRISPR backbone. The reaction was run for 20 cycles (37 °C 5 min/ 20 °C 5 min) and purified on the 25 nm cellulose membrane (MF-Millipore). The reaction was electroporated into Endura electrocompetent *E. coli* cells (Biosearch Technologies) to obtain the final plasmid pool. Typically, the library size was around 2 million clones.

To produce virus, 21x10^6^ HEK293T cells were seeded in T175 cell culture flasks in DMEM (Gibco) supplemented with 10% FBS (Gibco) and 1% penicillin/streptomycin (DMEM++). Cells were transfected the day after by preparing and mixing two reactions for 15 min at RT: the first consisted of 36 μg lentiviral library plasmid (pL.CRISPR or universal library), 21.6 μg psPAX2 plasmid and 7.2 μg pMD2.G plasmid (both gifts from Didier Trono, Addgene #12260 abd #12259, respectively) in 2 mL of DMEM and the second consisted of 120 μL polyethylenimine (PEI) in 2 mL of DMEM. After the reactions were mixed, 17 mL of DMEM++ was added and the total mix (21 mL) was added carefully on top of cells. The media was exchanged on the next day (15 mL) and the supernatant was collected 48 h thereafter. After collection, supernatant was passed through 0.45 μm filter units (Millex) and concentrated 150 times by ultracentrifugation (100,000 g for 2 h). Virus was re-suspended in stomach medium supplemented with 10 μM ROCK inhibitor Y-27632 (Sigma).

### Organoid transduction and selection

Organoids were expanded as described, harvested using mechanical disruption and pelleted by centrifugation. Chemical dissociation was then performed using TrypLE Express Enzyme (Gibco), by incubating organoids at 37 °C and checking dissociation process under the microscope, typically for 3-10 min. Dissociation was stopped using (DMEM)/F12 (Gibco) supplemented with 1x penicillin/streptomycin (Invitrogen), 1x Glutamax and 10 mM HEPES (DMEM+++). Cells were pelleted by centrifugation and re-suspended in complete stomach medium supplemented with 10 μM ROCK inhibitor Y-27632 (Sigma-Aldrich), 5 μg/mL protamine sulfate (Sigma-Aldrich) and concentrated virus re-suspended in stomach medium. The transduction mix was seeded in 48-well cell culture plate and centrifuged at 700 g for 1 h and then incubated at 37 °C for 4 h. The cells were harvested, seeded in Matrigel and supplemented with stomach media with 10 μM ROCK inhibitor. In the case of CRISPR-KO screen, organoids were selected with 2 μg/mL puromycin (Invivogen) for 6-7 days, starting on the third day after transduction. For CRISPRa screen, a dual selection was performed with puromycin (1.5 μg/mL) and blasticidin (10 μg/mL).

To determine the virus titer, CRISPR-KO and CRISPRa libraries were made in GFP backbone in addition to puromycin (CRISPR-KO) and blasticidin (CRISPRa) backbones. After GFP library infection, organoids were assessed for GFP expression by flow cytometry, and the titer achieving transduction efficiency of 20-25% was used for each of the screens.

### CRISPR-based hit validation pipeline

To validate the CRISPR screen hits, a GFP-based competition assay (Figure S1A) was performed in GC PDOs. Organoids were infected with lentiviral constructs (Cas9 or dCas9-VPR) and adjusted to obtain ∼ 1:1 mixture of genetically altered (GFP-positive, CRISPR-targeted) and wild-type (GFP-negative, unedited) cells. Mixed populations were split into two experimental arms: one treated with oxaliplatin (8 μM) for 72 h, and one left untreated. Surviving organoids were maintained and propagated, with GFP-positive fractions quantified weekly by flow cytometry for total of 4 treatment rounds.

To estimate GFP-positive population via flow cytometry, organoids were dissociated into single cells using TrypLE Express (Thermo Fisher). Cells were resuspended in 100–150 μL PBS supplemented with 0.04% BSA (Sigma). GFP fluorescence was measured on a FACSCalibur flow cytometer (BD Biosciences). Untagged wild-type cells were used as negative controls to define the gating strategy (Figure S1B). To gate the cells, forward versus side scatter (FSC-A/SSC-A) was used to identify live cell population and the doublets were then excluded using forward scatter height versus forward scatter area plot (FSC-H/FSC-A). Single cells were plotted in forward scatter area versus FITC area channel (FSC-A/FITC-A) and percentage of GFP-positive cells was determined. Data were analyzed with FlowJo Single Cell Analysis Software (BD Biosciences), and the relative share of GFP-positive cells was calculated for each sample. GFP enrichment in the drug-treated versus untreated arm was used as a readout of the resistance. To assess statistical significance, we compared each hit GFP enrichment over a time course to control (gene desert-targeting) gRNA via linear mixed effects model through the package lmerTest^45^ in R.

### gRNA efficiency quantification for the validated hits

For each of the validated hits, the efficiency of gRNA was estimated based on the type of alteration. In all cases, organoids were infected with single gRNA and selected with either puromycin (2 μg/mL, CRISPR-Cas9) or blasticidin (10 μg/mL, dCas9-VPR. For CRISPR-Cas9 screen hits, organoids were collected two weeks post-transduction, and gRNA editing efficiency was assessed by PCR amplification of the target locus, followed by bulk Sanger sequencing. Resulting chromatograms were analyzed using the Synthego ICE tool to estimate indel frequencies. For CRISPRa screen hits, total RNA was extracted from organoids using the RNeasy Mini Kit (Qiagen) with on-column DNase digestion (RNase-Free DNase Set, Qiagen). cDNA was synthesized from 0.5–2 μg RNA using the supertranscript IV Reverse Transcription Kit (Applied Biosystems, Thermo Fisher Scientific) with oligo-dT. Hit expression was estimated by quantitative PCR (qPCR) using the SYBR Green Master Mix (Thermo Fisher Scientific) on the real-time thermal cycler (CFX96; Bio-Rad). The same protocol was used to validate CRISPRa-based upregulation of *UBP1* in oxa-selected organoids. The sequences of all used qPCR primers and gRNAs are provided in Supplementary Table S9.

### Nucleic acid extraction and next-generation sequencing

RNA was extracted from PDOs using the RNeasy kit (Qiagen) and DNA using the DNA Blood Mini kit (Qiagen). For RNA-seq, poly(A)-selected libraries were prepared and sequenced on an Illumina NovaSeq 6000 (150 bp paired-end, ≥ 30M reads/sample). Exome sequencing libraries were prepared using the Agilent SureSelectXT Human All Exon V6 kit and sequenced at ≥ 80M paired-end reads/sample. For gRNA abundance estimation, genomic DNA was PCR-amplified in two steps: in the first reaction, 20 μg of genomic DNA was amplified with primers (F: 5′-GTAATAATTTCTTGGGTAGTTTGCA-3′; R: 5′-ATTGTGGATGAATACTGCCATTTG-3′) for 10 cycles. The product was then subjected to a second PCR with primers containing Illumina sequencing adapters (F: 5′-ACACTCTTTCCCTACACGACGCTCTTCCGATCTGGCTTTATATATCTTGTGGAAAGG-3′; R: 5′-GTGACTGGAGTTCAGACGTGTGCTCTTCCGATCTCAAGTTGATAACGGACTAGCC-3′) for 20 cycles. The final libraries were sequenced on Illumina NovaSeq, 250 bp paired-end.

### Exome sequencing data analysis

The reference genome was prepared using bwa index, and paired-end sequencing reads were aligned with BWA^46^ to human genome primary assembly hg38 under default settings. Resulting alignments were sorted, and PCR duplicates were identified and flagged using Picard MarkDuplicates. Mutations were identified using Varscan2^47^ and obtained variants were annotated with ANNOVAR^48^ and assessed for functional impact using SIFT^43^. Exome-seq datasets were also used to infer copy number changes using CNVkit^49^ To accomplish this, we utilized two normal stomach tissue-derived organoid lines as controls to build a virtual reference. Different tumor PDOs were then compared to the controls to estimate somatic copy number alterations specific to each line.

### CRISPR-screens analysis with MAGeCK

PCR-amplified gRNA libraries from both screens were analyzed with MAGeCK^50^ which was used both to generate gRNA count tables and to perform gene-level statistical analyses. In brief, MAGeCK quantifies gRNA abundance by mapping sequencing reads to the reference gRNA library, producing count tables for each sample. Counts were normalized for library size, and fold-changes were calculated by comparing endpoint to baseline samples. gRNAs were then ranked according to their log_2_ fold-changes.

To identify genes under positive selection, MAGeCK applies the robust rank aggregation (RRA) algorithm. This method calculates a positive selection score for each gene, reflecting the probability that its gRNAs are consistently enriched relative to the distribution of all library gRNAs. Each gene was represented by either four or five gRNAs; genes with multiple gRNAs clustering among the top-ranked positions were considered positively selected, resulting in lower positive selection scores. These positive selection scores as well as median log2 foldchanges for all genes were used in Figure 1E-F. Full results of the MAGeCK analyses are provided in the Supplementary Tables S6-7.

### RNA-seq analysis

Transcript quantification was performed with Kallisto^51^ collapsed to genes and imported to R via Tximport package^52^, where differential expression was analyzed with DESeq2^53^ Gene raw counts dataset from responders and non-responders (pre or post therapy) were obtained from the authors upon request^16^ and imported into R using Tximport package.

Prior to differential expression analysis, lowly expressed genes were filtered out to reduce noise and improve statistical power. Specifically, genes were required to have at least 100 raw counts in the smallest experimental group size to be retained.

The resulting filtered dataset was used to construct the final DESeq2 object. Differentially expressed gene lists were generated using the function contrast () from Deseq. Normalized expression counts were then obtained using the counts () function in DESeq2 with the argument normalized = TRUE, which accounts for library size factors and the lists of differentially expressed genes between parental and oxa-selected, and between parental and CRISPR-altered lines (UBP1 KO, HNF4A KO, SPDEF-upregulated, FOXQ1-upregulated) were generated. These normalized counts were subsequently used as input for gene set enrichment analysis (GSEA), ensuring that enrichment tests were performed on expression values corrected for sequencing depth.

### Gene set enrichment analysis

GSEA v4.0.3^24^ was applied using DESeq2-normalized counts generated in the previous step. Analysis was performed with 1000 geneset based permutations, gene set size 50–500 against three main molecular signatures databases: Gene ontology (biological processes), KEGG pathways, and hallmarks (v2023.1).

### TCGA copy number and survival analysis of *UBP1*

Copy number alteration (CNA) data for *UBP1* were downloaded from cBioPortal for Cancer Genomics (https://www.cbioportal.org). Copy number status was categorized as −1 (deletion), 0 (diploid), or +1 (amplification) according to GISTIC2.0 calls provided by TCGA. Cohorts were then defined as *UBP1*-deleted or *UBP1*-diploid based on these calls.

To integrate CNA with expression, RNA-seq data for TCGA-STAD (workflow type “STAR - Counts”) were downloaded from the Genomic Data Commons (GDC) using the TCGAbiolinks^54^ R package. Expression matrices were extracted from SummarizedExperiment objects, normalized, and mapped to TCGA barcodes. *UBP1* and *MAX* expression was retrieved and matched to CNA-defined cohorts. Next, these expression matrix TCGA cohorts stratified based on *UBP1* status were exported from R and used as an input to perform gene set enrichment analysis (GSEA). The analysis was performed with the same default parameters as described before, with the exception of permutations based on phenotype not geneset, since both groups had large enough size to conduct such an analysis as recommended by the GSEA documentation.

Statistical comparisons of survival between *UBP1*-deleted and diploid patient groups were performed using survival package in R via the function survfit^55^. Patients who previously received oxaliplatin were retrieved via cbioportal annotation of treatment and then overlapped with *UBP1*-deleted or diploid lists as identified previously in pooled rectal and colon adenocarcinoma cohorts READ+COAD. All analyses were carried out in R (v4.2.0).

### Oxaliplatin viability assays

For drug-response assays, organoids were fragmented into small clusters and seeded into 384-well plates (three biological replicates per condition; 21 wells each: six drug concentrations plus control, in triplicates). Each well contained 15 μL of Matrigel and 50 μL of gastric medium. Drug treatments were initiated on day 2 and refreshed on day 5, for a total of 6 days.

On day 7, cell viability was measured using CellTiter-Glo 3D (Promega). Thirty microliters of medium were replaced with reagent, plates were shaken for 25 minutes, and incubated for 5 minutes in the dark. Luminescence was recorded and used to generate dose–response curves for oxaliplatin using drm^56^ package in R.

### Organoid formation assay

To evaluate clonogenic capacity, PDOs were treated with drugs for 72 hours in 48-well plates (Corning), dissociated into single cells with TrypLE Express (Thermo Fisher Scientific), and counted manually. Single cells (3,000 per well) were seeded into six replicate wells of 96-well plates. After 7–8 days, organoids were imaged on a Celligo system (Nexcelom) and gated by organoid area around 2000 μm^2^. Formation efficiency was calculated as the percentage of single cells giving rise to organoids, normalized to vehicle-treated controls. Each condition was tested in three independent biological replicates for all PDOs tested. For DD109 PDOs, we used IC_30_ oxaliplatin for each line, 3 μM for parental and UBP1 KO PDOs, and 15 μM for oxa-selected, while 4EGI 40 μM was used for all lines. For OO99 PDOs, we used the same concentration of oxaliplatin for both lines (6.5 μM) and 33 μM 4EGI for both pre-therapy and post therapy PDOs.

### Organoid cytotoxicity assay

To quantify treatment-induced cell death, PDOs were dissociated into single cells and reseeded in 25μL Matrigel domes in 96 well plate (Greiner, Cat No 655866). After 4 days, formed organoids were treated with indicated drugs and combinations. After 3 days, treated organoids were incubated for 1 h with Hoechst 33342 (Invitrogen) and propidium iodide (PI, Sigma) for a final concentration of 10 μg/ml each to stain total and dead nuclei, respectively. Hoechst-only control wells were used to estimate background PI signal. Bortezomib (10 μM) was used as a positive control and DSMO was used as a negative control. Raw percentages of PI positive cells were normalized via min-max normalization between DMSO and Bortezomib wells to obtain percentage of PI-positive organoids used to draw plots in Figure 5 F-H.

### Image based quantification of cytotoxicity in PDOs

Image acquisition was performed on a Yokogawa CV7000 high-content imaging system. Brightfield (Trasmitted light), Hoechst (Excitation laser 405 nm, Emission bandpass filter 445/45) and Propidium Iodide (PI, Excitation laser 561, Emission bandpass filter 600/37) channels were acquired in live acquisition mode at 37° C and 5% CO2 with an Olympus UPLSAPO 10 X objective (NA 0.4), sampling 7 fields per well across three Z-planes, optically slicing the imaging volume every 30 mm. Nuclei were segmented from the Hoechst channel using StarDist^57^, enabling reliable and accurate nuclear segmentation in the organoid context. The segmentation labels were then imported into CellProfiler (version 4.2.6; https://cellprofiler.org/previous-releases) together with the images from all channels, allowing nuclear features to be placed within the organoid architecture, to quantify PI signal intensity for each nucleus and to generate image overlays as png files to visually monitor segmentation performance.

Both the StarDist-based segmentation and subsequent CellProfiler analyses were executed on the MPI-CBG computing cluster using Python scripts to enable parallelization. Quantitative results were exported as tabular data and further processed in KNIME (version 4.7.8; https://www.knime.com/downloads/previous). The proportion of PI-positive cells was determined by applying a threshold derived from the distribution of PI fluorescence intensities in negative control cells (cells not stained with PI) thereby classifying nuclei as PI-positive or -negative, minimizing technical false positives.

### Polysome profiling and analysis

PDOs were split twice every 48 h before collection. On the day of collection, PDOs were pretreated with cycloheximide (100 μg/mL, Sigma) for 30 min, harvested with cell recovery solution (Corning) containing 100 μg/mL cycloheximide, and pelleted. Pellets were stored at −80 °C. For lysis, frozen pellets were thawed on ice and resuspended in polysome lysis buffer (20 mM Tris-HCl pH 7.4, 150 mM NaCl, 5 mM MgClC, 1 mM DTT, 100 μg/mL cycloheximide, 1% Triton X-100, and 24 U Turbo DNase; Invitrogen). Lysates were incubated on ice for 15 min, sheared 10 times through a 25G needle, and clarified by centrifugation (10 min, 4 °C). RNA concentration was quantified by absorbance at 260 nm (NanoDrop, Thermo Fisher), and equal input was used for gradient loading 2.8 A260 units for DD109 as well as OO99 experiment. Sucrose gradients were prepared fresh in UltraClear tubes (14 × 95 mm, Beckman Coulter) by overlaying 6 mL of 10% sucrose solution on top of 6 mL of 50% sucrose solution, both prepared in polysome buffer. Gradients were linearized (5–45%) using a Gradient Master (Biocomp). Lysates were layered onto gradients and centrifuged at 36,000 rpm for 3 h at 4 °C in an SW40 rotor (Beckman). Gradients were fractionated on an ÄKTA Pure chromatography system (Cytiva) equipped with a UV detector set to 260 nm and a peristaltic pump.

Polysome profiles were recorded and aligned horizontally to center around the monosome peak, as well as baseline corrected by subtracting the lowest point of each profile from all values and add +1 pseudocount to avoid zero values using R custom code. Aligned profiles were subsequently used to quantify different areas under curves that corresponds to either polysomes (first 4 peaks to the left of profile) or monosomes (central peak) through a custom R-based Shiny app. The resulting values were used to calculate polysome to monosome ratio corresponding to each profile. The output quantification is attached as supplementary material 1 and 2.

### Western blotting

PDOs were collected mechanically and lysed on ice for 1 hour in RIPA buffer (50 mM Tris pH 8.0, 0.1% SDS, 0.5% sodium deoxycholate, 150 mM sodium chloride, 1% Triton X-100) supplemented with the Halt protease and phosphatase inhibitor cocktail (Thermo Fisher Scientific, #78440). Protein concentrations were measured using the Pierce BCA protein assay kit (Thermo Fisher Scientific, #23225). Lysates (10 μg protein) were denatured at 85°C for 3 min, separated on 4–12% Bis-Tris gels (NuPAGE, Thermo Fisher Scientific), and transferred to nitrocellulose membranes (Amersham Protran, GE Healthcare). Membranes were blocked with 5% non-fat milk in PBS-T and incubated overnight at 4°C with antibodies against 4E-BP1 (#9452, CST, 1:1000), phospho-4E-BP1 (S65; #9451, CST, 1:1000), GAPDH (TA302944, Origene, 1:10000), or β-tubulin (1:1000, Santa Cruz Biotechnology, sc-166729). The next day, membranes were washed three times for 10 min each in 5% milk PBS-T and then incubated for 40 min at room temperature with secondary antibodies (IRDye 800CW and IRDye 680LT, both LI-COR Odyssey, 1:13000). Bands were visualized with the LI-COR Odyssey imaging system. As a molecular weight standard, Spectra Multicolor Broad Range Protein Ladder (Fermentas) and MagicMark XP Western Protein Standard (ThermoFisher) were used.

### Immunofluorescence imaging-based analysis of nucleolar integrity

Parental, UBP1 KO, and Oxaliplatin-selected PDOs, treated with either vehicle or oxaliplatin (4 μM), along with a positive control consisting of the parental line treated with rapamycin (20 μM), were prepared for imaging as previously described^12^. Imaging was performed using an Andor Spinning Disk System equipped with a 60×/1.3 NA silicon oil objective and an additional 2× magnification. All images were deconvolved using Huygens. Image analysis was conducted through an initial preprocessing step applying a Gaussian denoising filter (1 μm diameter) to both nuclei (DAPI) and protein channels. Nuclei segmentation was performed using a pretrained Cellpose model (nuclei diameter = 8 μm, cell probability threshold = 0.0, flow threshold = 0.4). Protein signal segmentation was achieved by applying a blob-finder algorithm to the protein channel (blob diameter = 0.5 μm, probability threshold = 20%). Colocalization analysis was then carried out to identify protein segments fully overlapping with nuclei masks (100% overlap). The volumes of colocalized protein segments were normalized to the volume of the corresponding nuclei to generate a “Ratio_volumes” metric, which was utilized for downstream plotting and analysis. Ratios in oxaliplatin-treated samples were normalized to the median value of corresponding untreated samples. Statistical comparison was performed by Wilcoxon test.

### Chromatin immunoprecipitation (ChIP)

To perform chromatin immunoprecipitation (ChIP), we first generated an N-terminal GFP fusion to UBP1. For this purpose, UBP1 was amplified from cDNA isolated from DD109 and cloned via MluI site into the universal plasmid^12^ (primer sequences given in Supplementary Table S9). DD109 organoids were infected with lentivirus carrying GFP-UBP1 fusion at low MOI to prevent high overexpression. Furthermore, infected cells were sorted by FACS for low GFP expression. The resulting organoids were used for the ChIP-seq experiment.

Chromatin immunoprecipitation was performed as previously described with minor modifications^12^. After organoid dissociation, six million cells were fixed with 1% (v/v) formaldehyde (methanol free) at room temperature for 10 min. Crosslinked chromatin was prepared and resuspended in 1ml shearing buffer D3 (Covaris, truChIP® Chromatin Shearing Kits). Chromatin was sheared to an average size of 300-500 bp by a Covaris sonicator with the setting Duty factor 5, 200 cycles per burst for 8 minutes. ChIP assays were performed as previously described using 30 ug polyclonal goat anti-GFP antibody that was generated in Max Planck Institute of Molecular Cell Biology and Genetics^58^. Following overnight incubation with antibody, genomic DNA was immunoprecipated with protein G Sepharose bead (GE healthcare, 17-0618-01). Reverse cross-linking was performed in the presence of Proteinase K at 65 °C overnight. After treatment with RNase A, Genomic DNA was cleaned up using Genejet PCR purification (Thermo Fisher Scientific, K0701) and quantified by Qubit™ dsDNA HS Assay Kit (Thermo Fisher Scientific, Q32854). Around 58 ng of DNA was used for the Illumina sequencing.

### ChIP-seq analysis

Sequencing reads were aligned to the human reference genome (GRCh38 primary assembly) using BWA-MEM v0.7.17^46^. The genome was indexed with bwa index, and paired-end reads were mapped with default parameters. Aligned reads were sorted and PCR duplicates were marked using Picard MarkDuplicates.

Peak calling was performed with MACS3 v3.0.0a7^59^ in paired-end mode, using matched input DNA as control. A q-value threshold of 0.01 was applied, with a bandwidth of 300 bp and model fold range of 5–50. The effective genome size was set to 2.7 × 10C (human). MACS3 estimated the average fragment length at 292 bp. For the UBP1_GFP ChIP sample, 38.2 million fragments were sequenced, of which ∼29.8 million remained after filtering. For the input control, 38.9 million fragments were sequenced, of which ∼29.3 million passed filtering. Called peaks were annotated with CHIPseeker and used to infer feature distribution of called peaks as well as its distribution around transcription start site. The annotated peaks from CHIPseeker are provided in Supplementary Table S8.

Signal tracks were generated from the treatment and control pileup bedGraph files using bedGraphToBigWig (UCSC Genome Browser utilities) with GRCh3 8 chromosome size reference. Visualization was performed with pyGenomeTracks^60^ using custom track configuration files.

### Statistical analysis

For gene expression comparisons, we used Deseq’s generalized linear models to compare different conditions with minimum of three biological replicates per condition. Deseq uses Wald test and reports p value along with Benjamini–Hochberg corrected p value which we use to indicate statistical significance. For comparisons with no replicates, i.e. pooling different patients in one condition, we use Wald test’s p value as measurement of statistical significance due to the high level of inter patient heterogeneity.

For organoid formation assay, we used paired t-test per replica and resulted p value was used as measurement of significance. For cytotoxicity assay, we use Student’s t-test to indicate statistical significance between the three replicates of each condition.

## Data Availability

Data and analytical methods are available to other researchers upon request to the corresponding authors. Patient-derived organoid models can be made available upon approval of the local ethics committee.

