## supplementary figures for "Loss of *UBP1* drives oxaliplatin resistance through a targetable dependency on translation initiation"

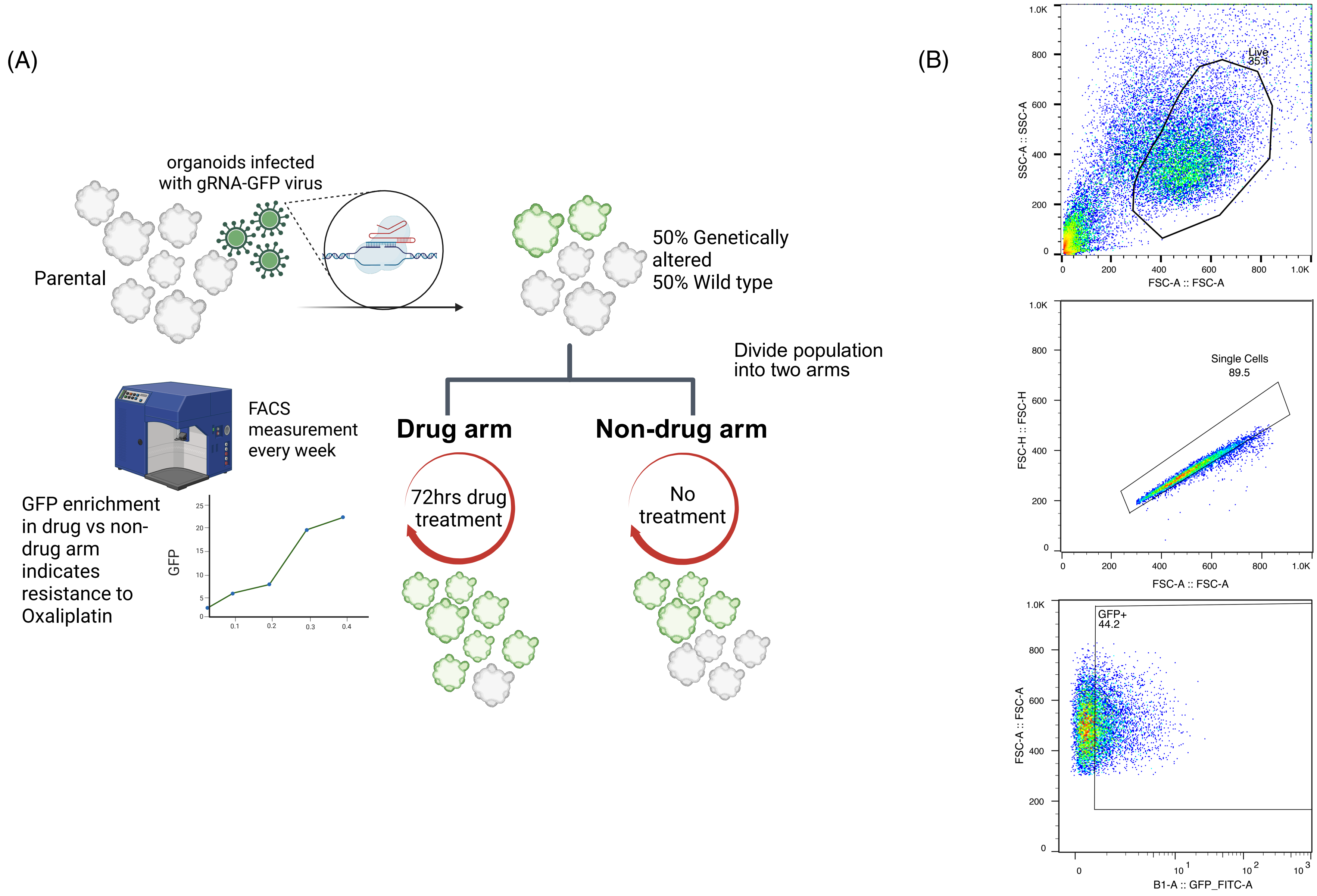


**Figure S1. CRISPR-based hit validation (organoid competition) pipeline in gastric cancer PDOs. (A)** The schematics of the pipeline. Crucial steps are highlighted. **(B)** Flow cytometry gating strategy to define GFP-positive (CRISPR-edited) cell population.

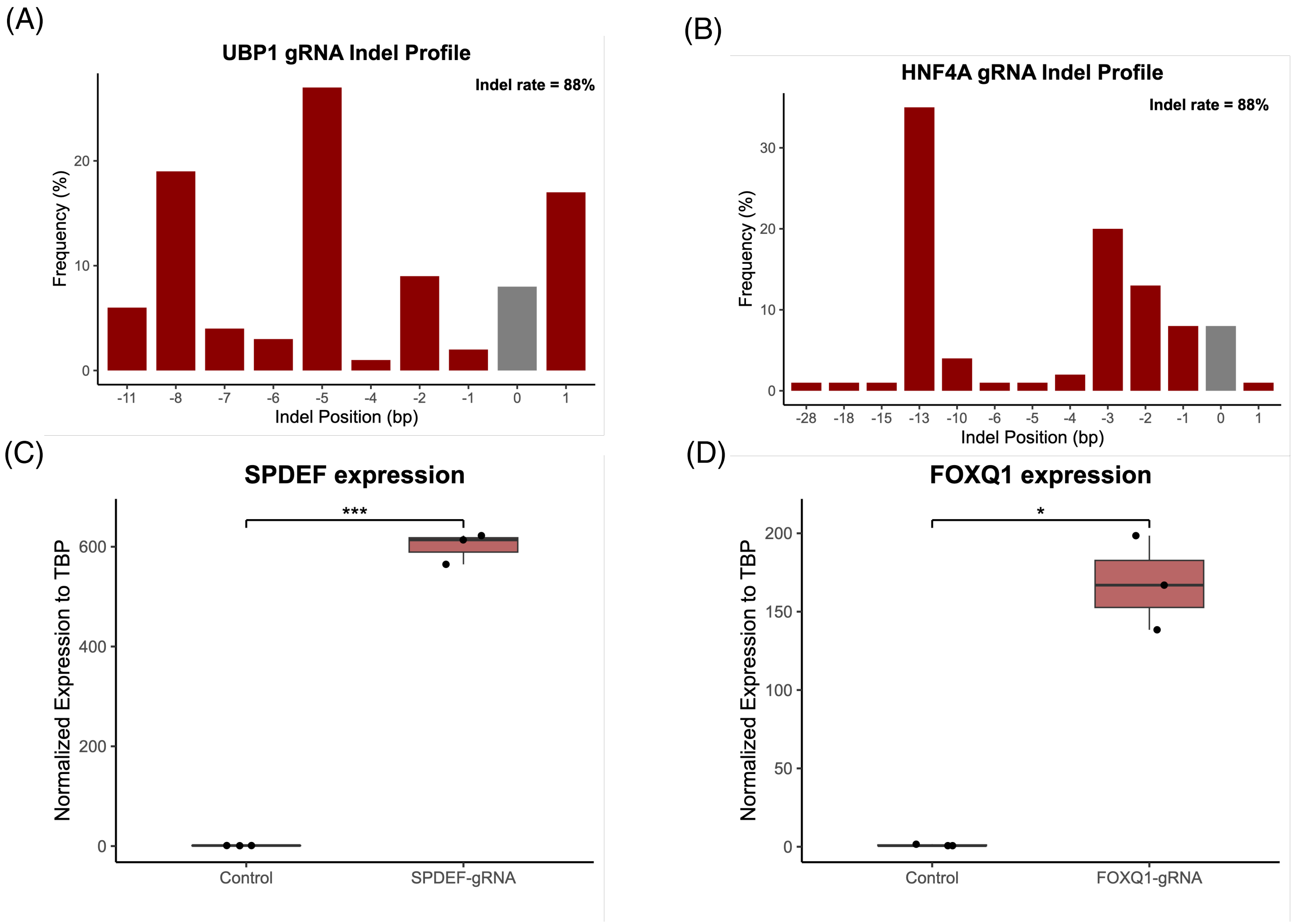


**Figure S2. gRNA validation of the screen hits.** Indel profiles are depicted for the knockout CRISPR-Cas9 screen hits (**A and B**). Quantitative PCR results for the upregulation screen hits are shown in panels **(C)** and **(D)**. Statistical comparison was performed using t-test. ***$p < 0.001$ and *$p < 0.05$.

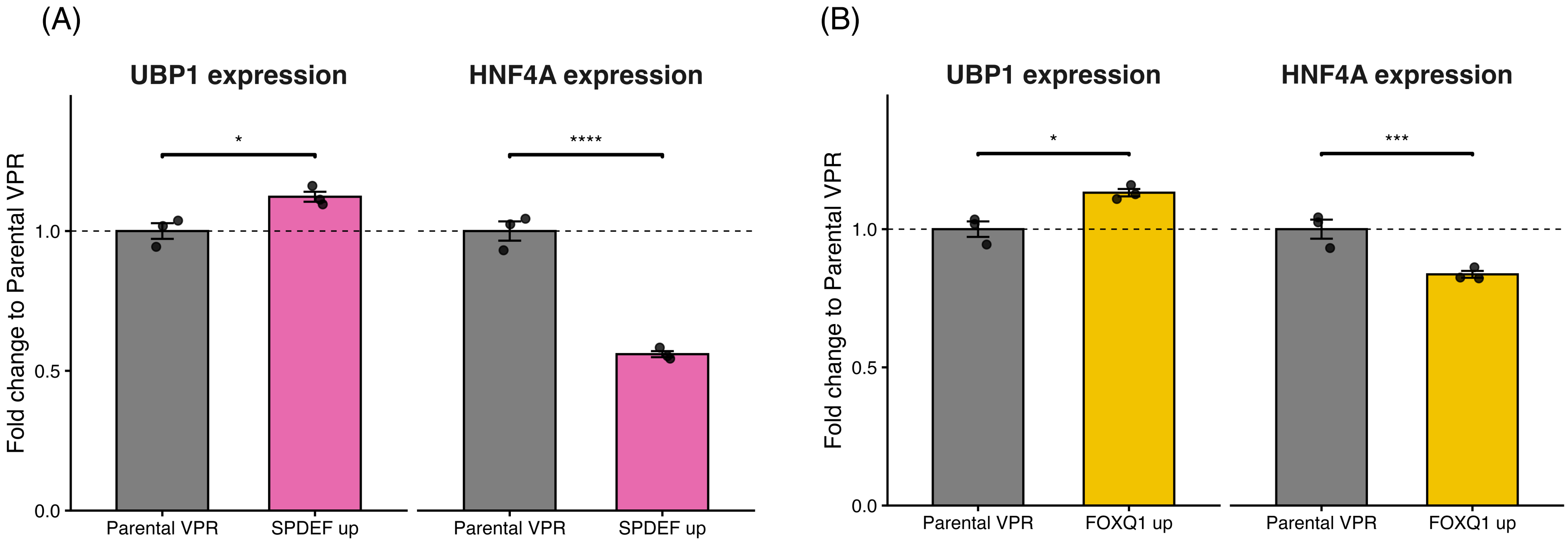


**Figure S3. Upregulation of *SPDEF* and *FOXQ1* does not recapitulate changes in all screen hits. (A)** *UBP1* and *HNF4A* expression in SPDEF-upregulated organoids compared to parental VPR organoids. **(B)** *UBP1* and *HNF4A* expression in FOXQ1-upregulated organoids compared to parental VPR organoids. All statistical comparisons were performed by Deseq's Wald tests (*$p < 0.05$, ***$p < 0.001$, and ****$p < 0.0001$).

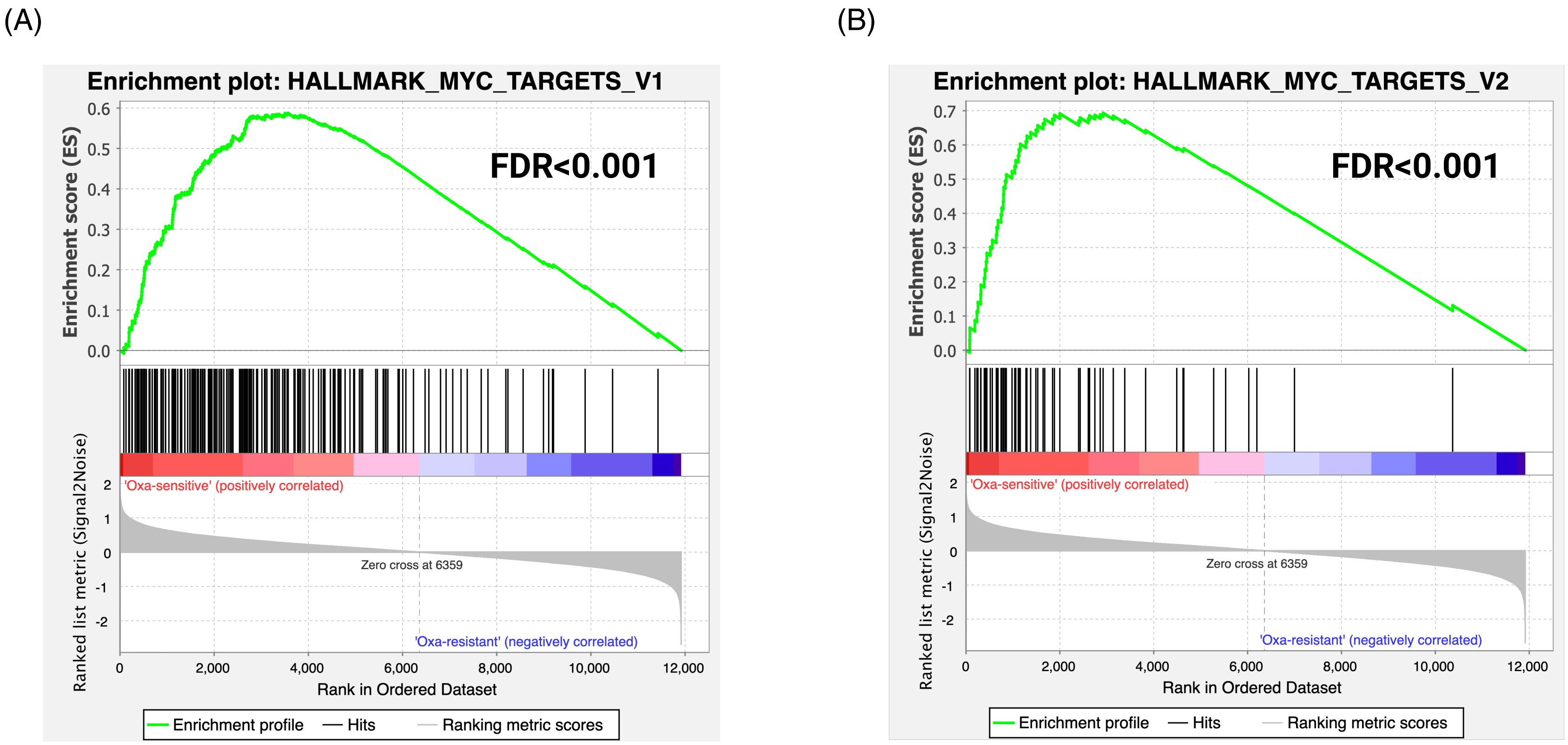


**Figure S4. GSEA enrichment plots comparing oxaliplatin-sensitive versus oxaliplatin-resistant patient-derived organoids from the OPPOSITE co-clinical trial.** Enrichment is shown for two gene sets: the Hallmark “MYC Targets V1” **(A)** and “MYC Targets V2” **(B)** signatures.

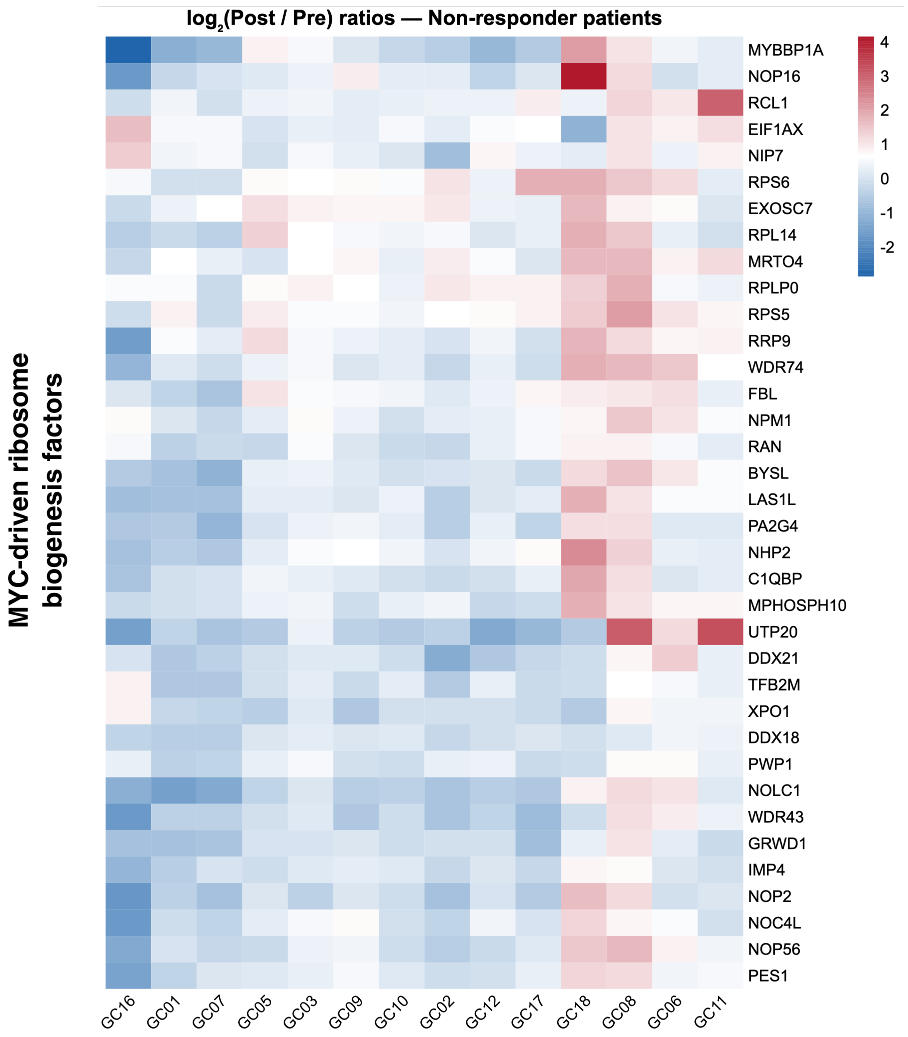


**Figure S5. Expression of MYC-driven ribosome biogenesis factors in post-treatment vs. pre-treatment tumors.** Heatmap of $log_2$(post/pre) treatment endoscopic biopsy sample expression ratios for a curated MYC-driven ribosome biogenesis factor signature in the same non-responder cohort (columns = individual patients; rows = genes).

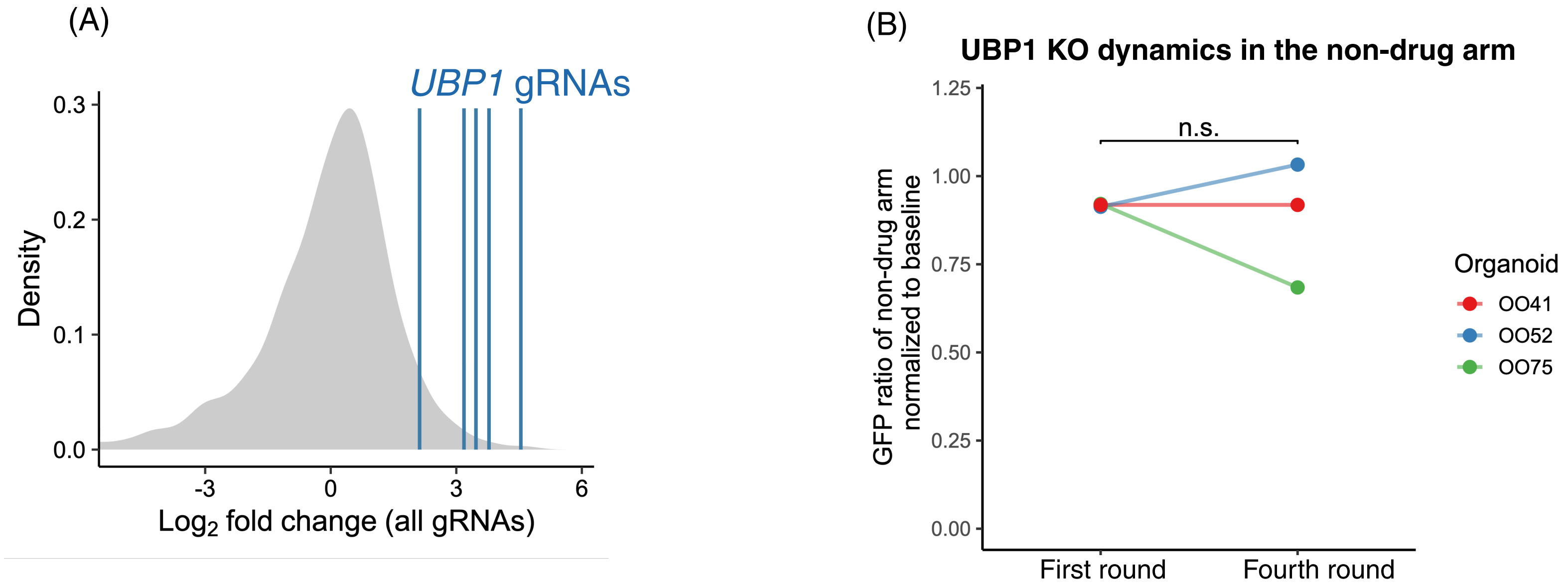


**Figure S6. Effects of *UBP1* gRNAs in the presence and absence of oxaliplatin. (A)** Rankings of *UBP1* gRNAs among all screen gRNAs ordered by $\log_2$ fold-change. **(B)** UBP1 gRNA effects in the absence of oxaliplatin in three PDOs from the OPPOSITE co-clinical trial. Statistical comparison was performed using paired Student's t-test. n.s. - not significant.

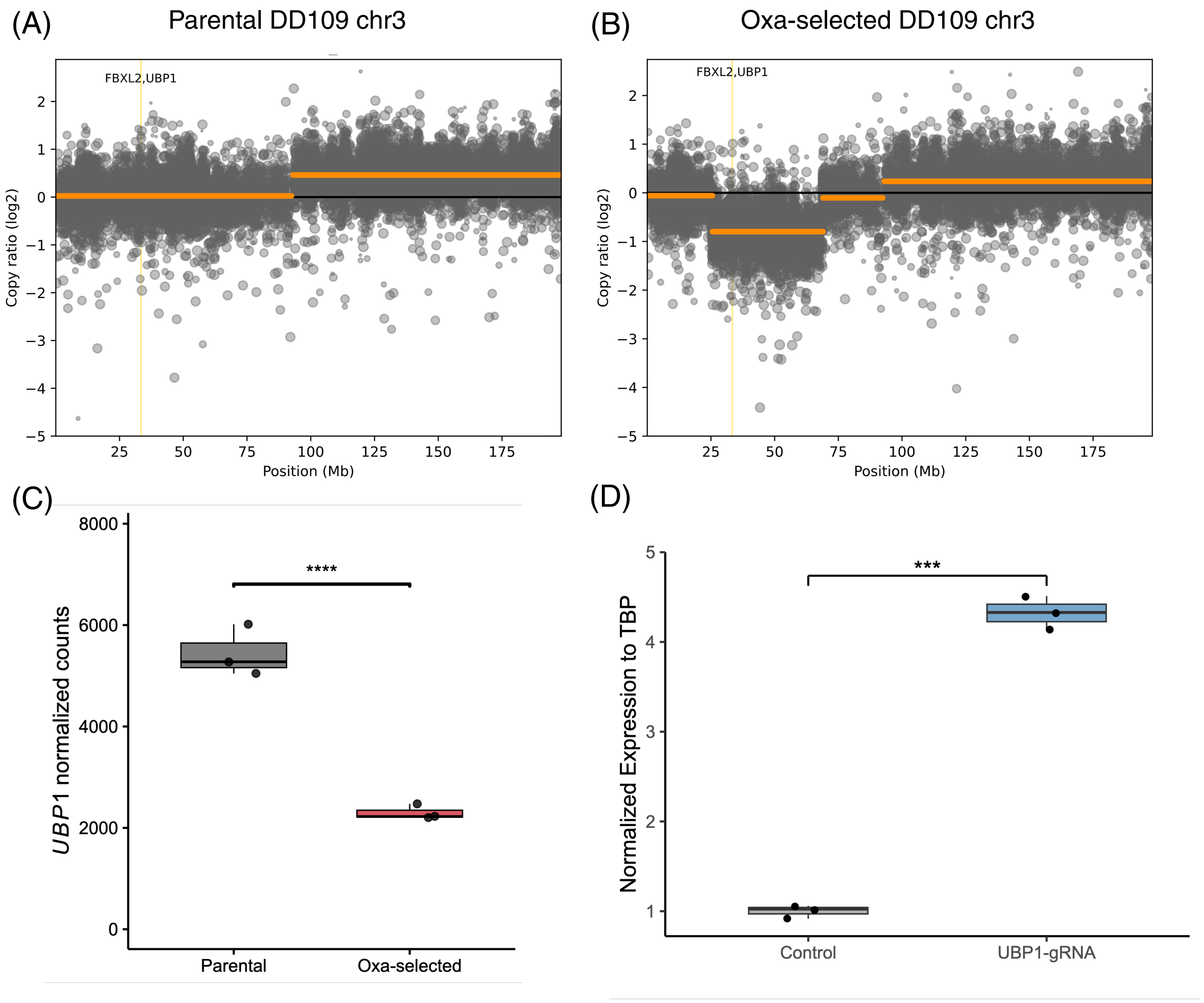


**Figure S7. UBP1 genomics and transcriptomics.** Copy ratios of chr3 in parental **(A)** and oxa-selected PDO **(B)** are depicted. **(C)** Boxplot of *UBP1* normalized counts in parental versus oxa-selected organoids, with Benjamini–Hochberg–adjusted p-value indicated above the comparison. ****p < 0.0001. **(D)** Boxplot of quantitative PCR results for *UBP1* upregulation via CRISPRa system. The statistical comparison was performed using Student's t-test, ***p < 0.001.

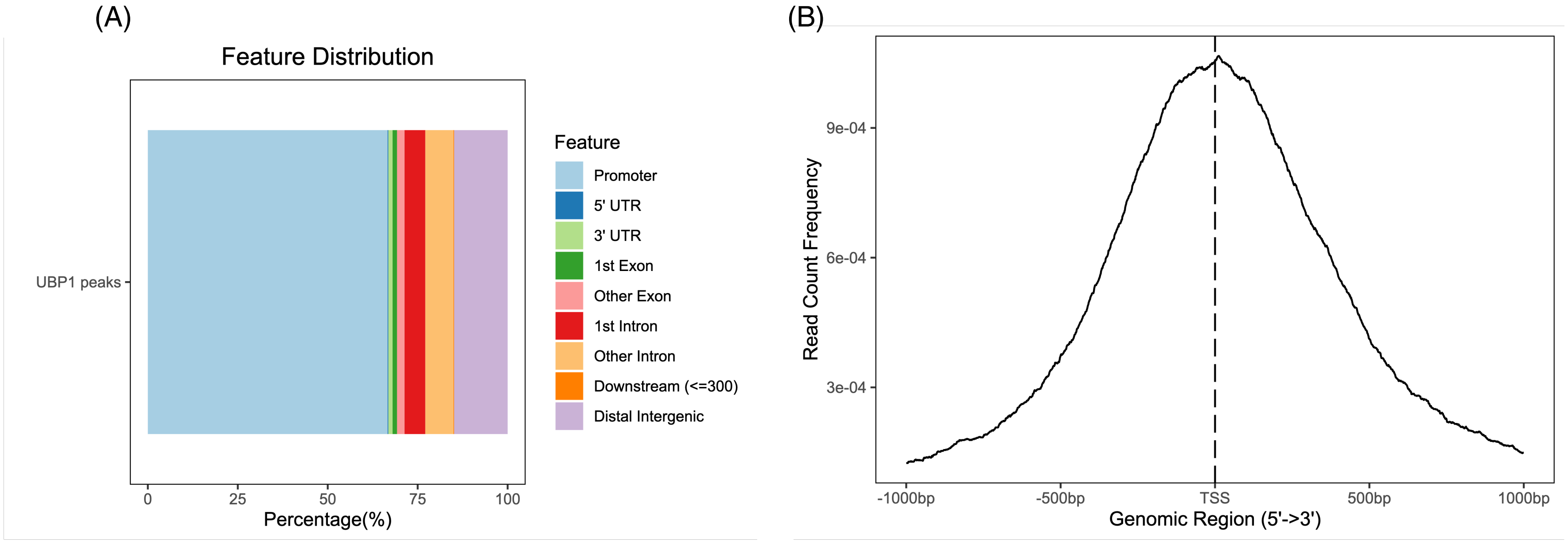


**Figure S8. UBP1 CHIP-seq peaks annotation and distribution around TSS. (A)** Bar plot of feature distributions for UBP1 peaks, showing the percentage of peaks located in different gene elements. **(B)** CHIP-seq read count frequency across ±1 kb around the transcription start site (TSS).

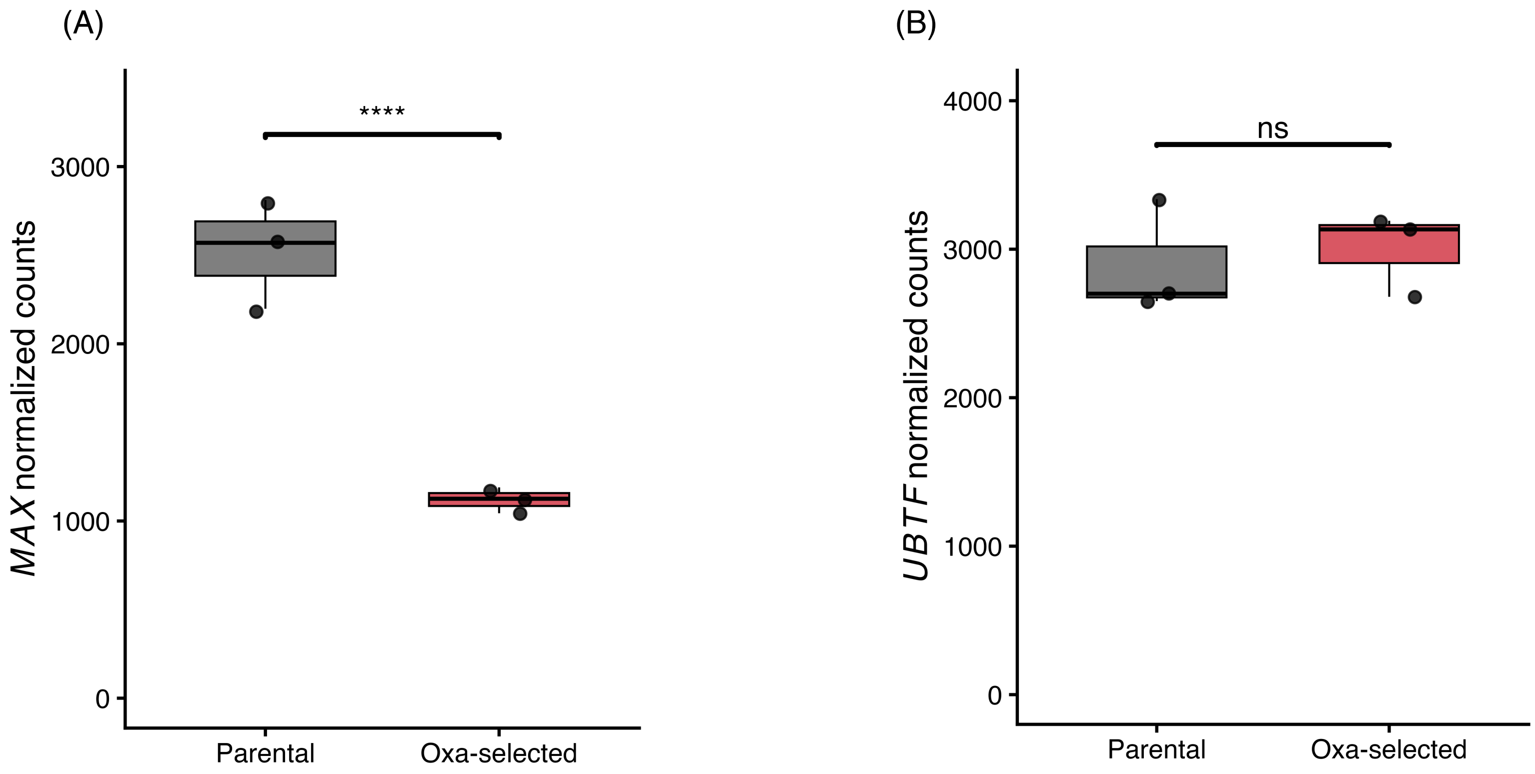


**Figure S9. *MAX* and *UBTF* expression in Oxa-selected organoids. (A)** Boxplot of *MAX* normalized counts in parental versus oxa-selected organoids, with Benjamini–Hochberg–adjusted p-value indicated above the comparison. ****p < 0.0001. **(B)** Boxplot of *UBTF* normalized counts in parental versus oxa-selected organoids, with Benjamini–Hochberg–adjusted p-value indicated above the comparison. ns~non-significant.

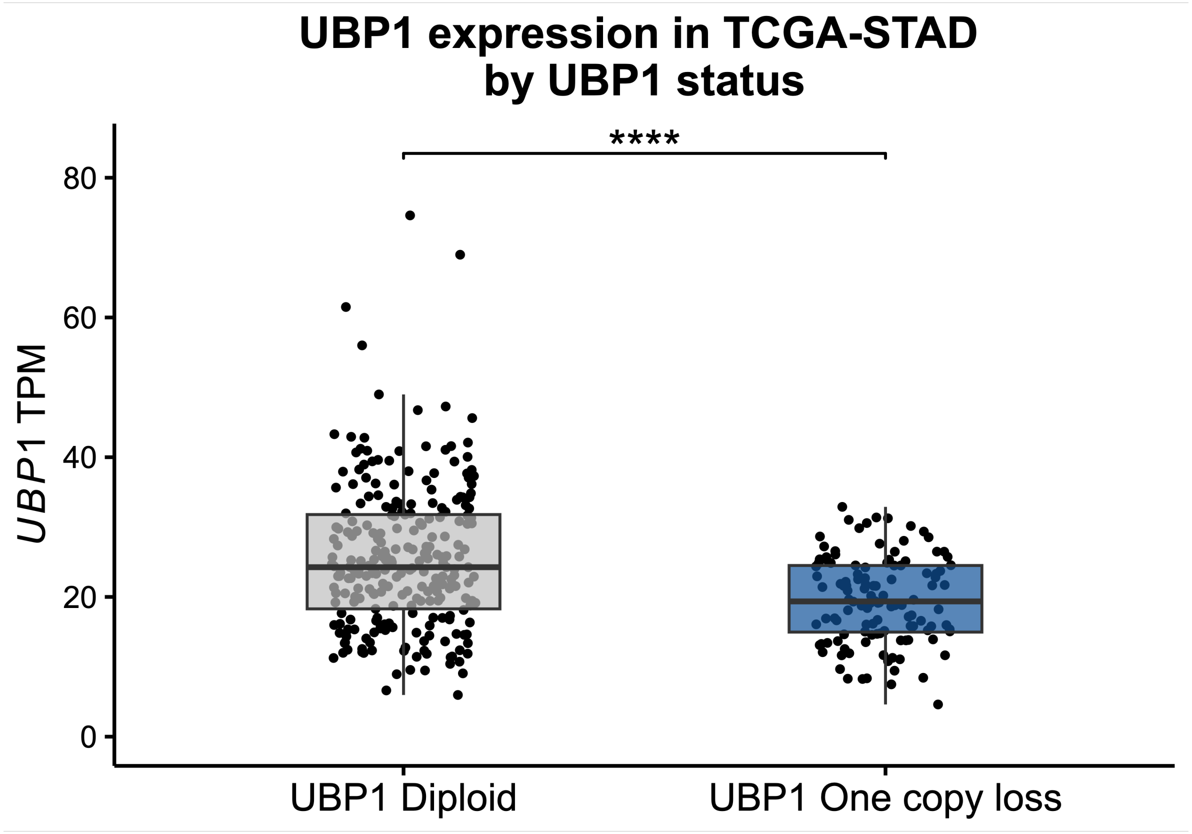


**Figure S10. Boxplot of *UBP1* mRNA expression (TPM) in TCGA stomach adenocarcinoma (STAD) samples stratified by *UBP1* copy-number status (diploid vs. one-copy loss).** Each point represents one sample. Statistical difference was calculated by t-test for effect size measured by Cohen's d. p-value (****p < 0.0001) and Cohen's d=0.652. Cohen's d>0.5 indicates statistically significant medium to large effect.

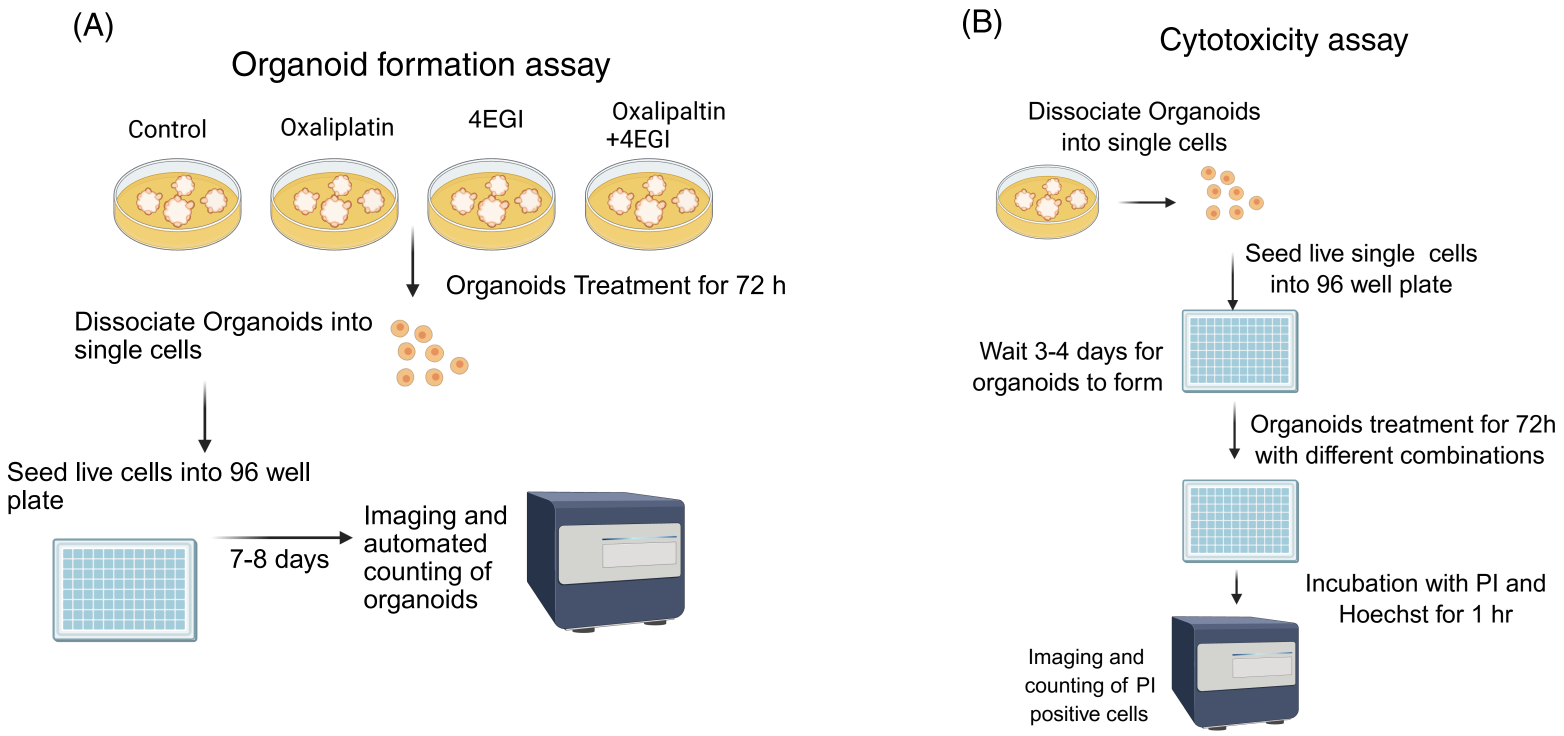


**Figure S11. Workflows for treatment effect estimations. (A)** Organoid formation assay workflow. **(B)** Cytotoxicity assay workflow. Crucial steps are highlighted.

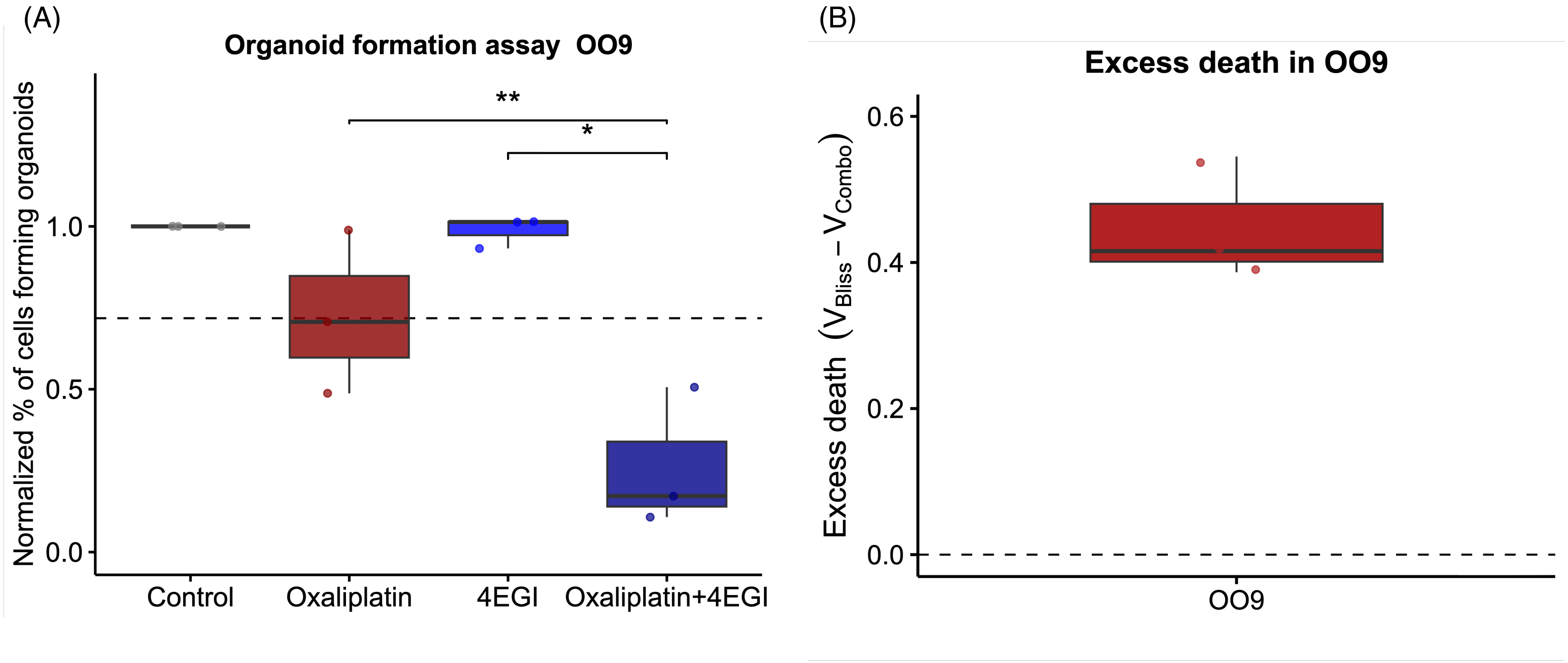


**Figure S12. Reversing oxaliplatin resistance in OO9. (A)** Organoid formation assays in the most resistant PDO from the co-clinical OPPOSITE trial treated with control treatment (DMSO), Oxaliplatin (20 µM), 4EGI (40 µM), and a combination of both. Organoid-forming efficiency is normalized to the DMSO control. Dashed line represents expected additive effect under the Bliss independence model. **(B)** Excess death calculated as the difference between expected and observed organoid formation capacity in OO9 PDO model under the Bliss independence model. Horizontal bars denote paired student t-test comparisons (*p < 0.05, **p < 0.01).

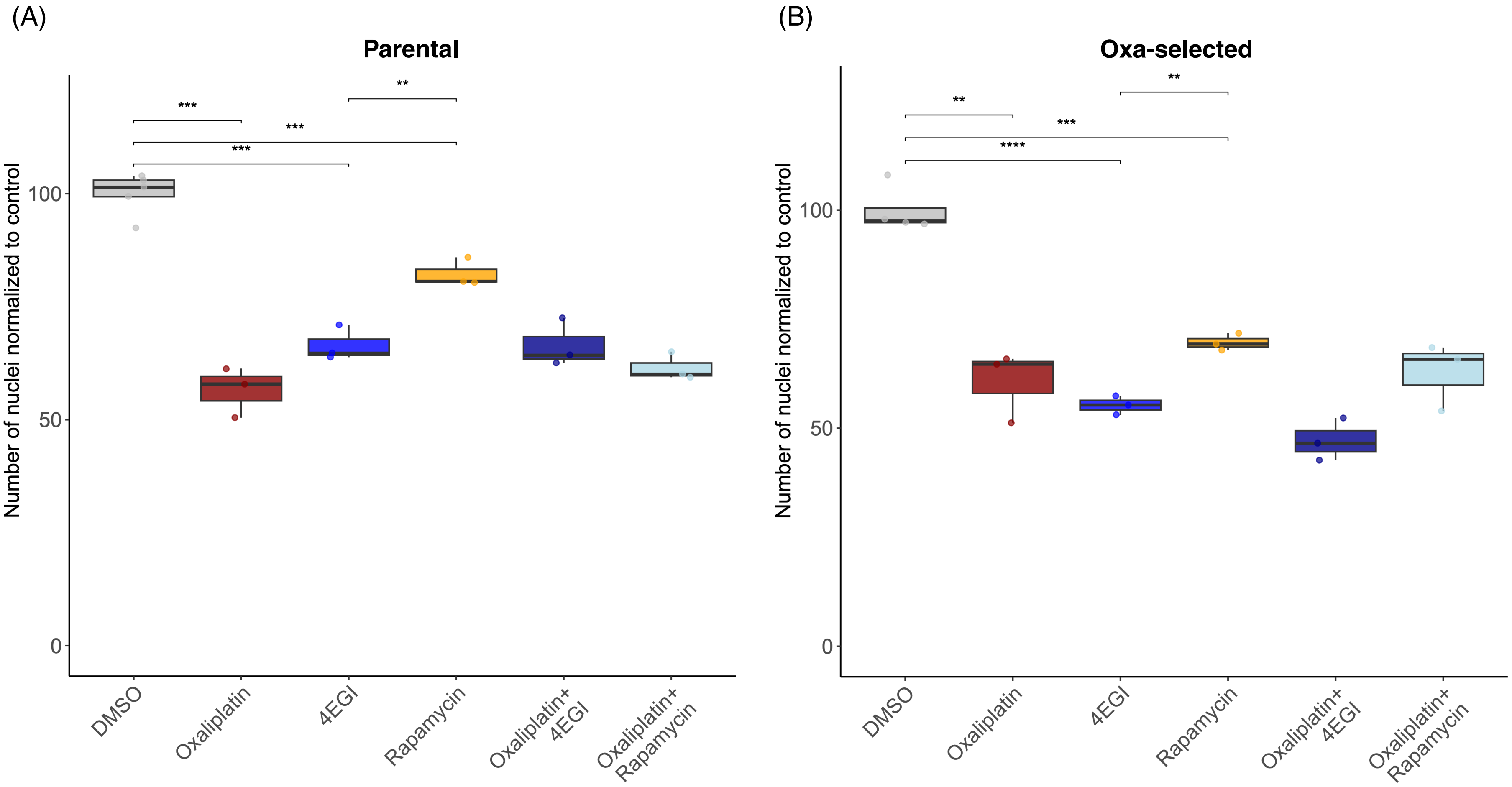


**Figure S13. Effects of treatments on organoid growth in the cytotoxicity assay. (A)** Boxplot showing the number of nuclei per well (normalized to the DMSO control) in parental organoids after 72 h treatment with single agents; oxaliplatin (10 µM), 4EGI (50 µM), rapamycin (20 µM) or the indicated combinations (oxaliplatin + 4EGI, oxaliplatin + rapamycin). Each box depicts the median, interquartile range and full range of three replicates; individual points are shown. Horizontal bars denote pairwise Student's t-test comparisons (*p < 0.05, **p < 0.01, ***p < 0.001, ****p < 0.0001; ns = not significant). **(B)** Same analysis as in **(A)**, but performed in oxa-selected PDO.

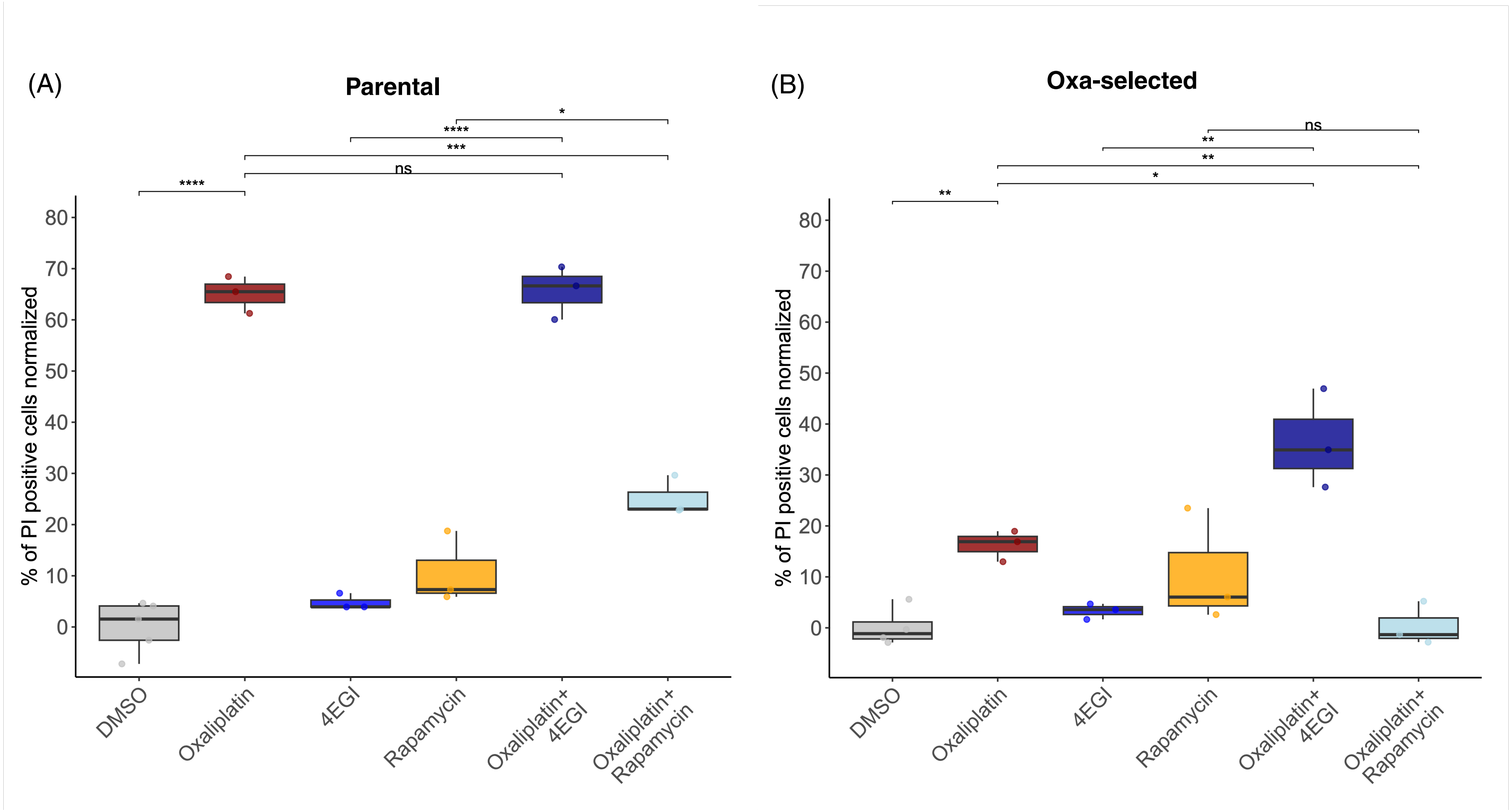


**Figure S14. Effects of treatments on cell death in the cytotoxicity assay. (A)** Boxplot showing the percentage of propidium iodide-positive nuclei out of total nuclei (min-max normalized to the DMSO-Bortezomib, 10 µM) in parental organoids after 72 h treatment with single agents; oxaliplatin (10 µM), 4EGI (50 µM), rapamycin (20 µM) or the indicated combinations (oxaliplatin + 4EGI, oxaliplatin + rapamycin). Each box depicts the median, interquartile range and full range of three replicates; individual points are shown. Horizontal bars denote pairwise Student's t-test comparisons (*p < 0.05, **p < 0.01, ***p < 0.001, ****p < 0.0001; ns = not significant). **(B)** Same analysis as in **(A)**, but performed in oxa-selected PDO.

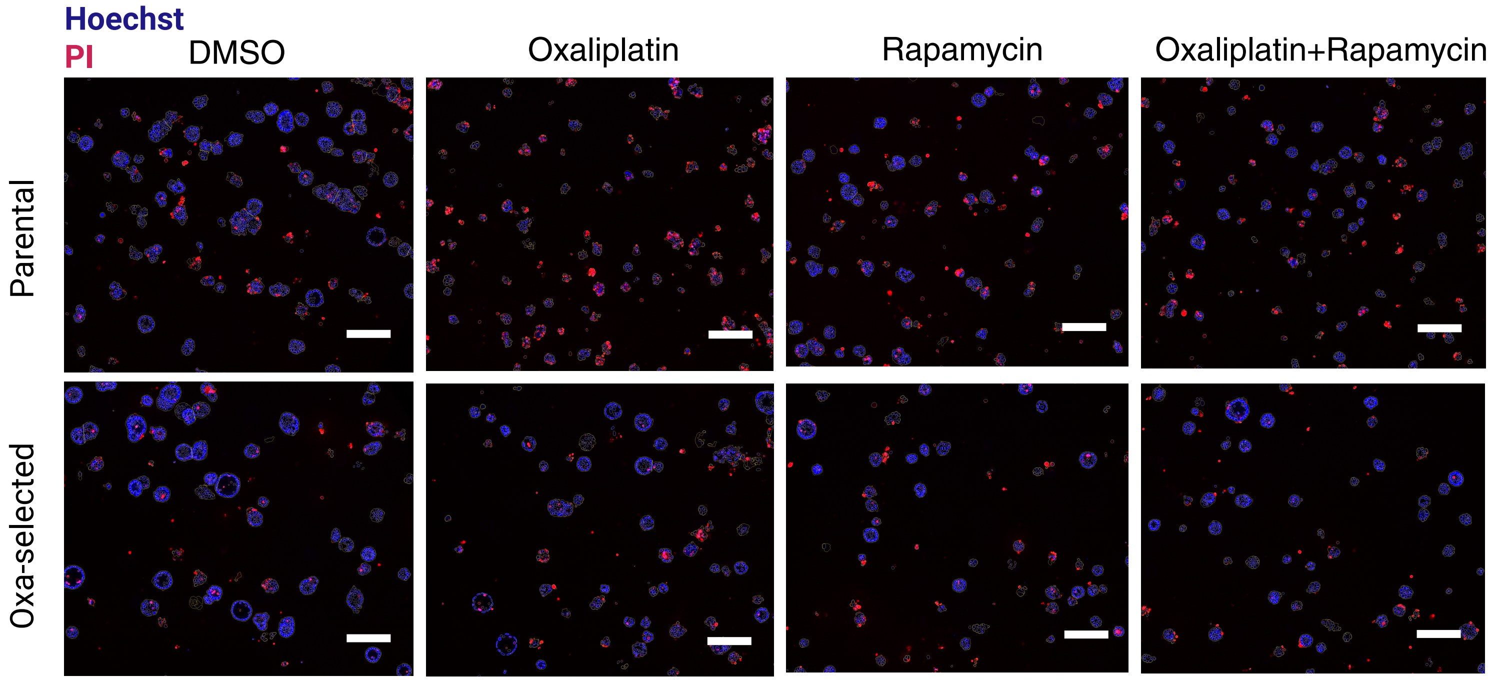


**Figure S15. Representative images of effects of oxaliplatin and rapamycin treatments on cell death in the apoptosis assay.** Scale bars represent 200 µm.
